# Starvation modulates associative short-term memory of *Drosophila* in a task-dependent manner

**DOI:** 10.64898/2026.06.09.729932

**Authors:** Edanur Sen, Svea Königsmann, Mozhdeh Besharatifar, Anna Ciuraszkiewicz, Sevval Demirci, Ayse Irem Güler, Thomas Niewalda, Michael Schleyer, Michael Thane, Christian König, Juliane Thoener, Bertram Gerber

## Abstract

It is widely believed that starvation favours the processing of food-related cues, a notion here called the ’adaptive specificity hypothesis’. Indeed, in *Drosophila melanogaster* starvation is required for appetitive odour-sugar but not for aversive odour-shock memory. Results from Gruber et al. (2013) and Meschi et al. (2024), however, suggest that starvation improves aversive short-term memory, too, challenging this hypothesis. We survey how starvation affects *Drosophila* associative olfactory short-term memory across 26 learning tasks. These tasks differ in the reinforcers and the amount of training, the life stage of the animals, in the predictive structure and associative timing of the task, in whether memory is expressed as an increase or decrease in odour preference, and in whether the learned behaviour is motivated by obtaining reward or avoiding/ escaping punishment.

In adult flies, an improvement was observed for appetitive odour-sugar memories, whereas all tasks yielding aversive memory were unaffected. Strikingly, ’appetitive’ tasks that are not sugar-related, namely odour-shock extinction learning and punishment-relief associations, were either unaffected or even impaired, supporting the ’adaptive specificity hypothesis’. In contrast, in 5-day-old larvae sugar-related appetitive associations were compromised, and the same was observed, to varying degrees, in larvae starved one day earlier and for aversive quinine associations, challenging the ’adaptive specificity hypothesis’. Furthermore, we observed starvation-induced changes in locomotion and preference for a subset of the cues used in our study.

Our results defy a simplistic interpretation in terms of the ’adaptive specificity hypothesis’ and call for case-by-case analyses of how starvation affects learning and behaviour.

## Introduction

Animals under starvation develop a syndrome of metabolic, physiological and behavioural changes aimed at restoring nutritional homeostasis, with ’hunger’ as the psychological corollary of this syndrome. As under starvation food is absent but sought-for, processing and learning about food-predicting cues is of particular biological significance. In this context we ask how starvation affects memory in *Drosophila melanogaster*. These animals are simple enough to be tractable and complex enough to be interesting for such an analysis (Sgammeglia and Sprecher, 2022; Suarez-Grimalt et al., 2024). Moreover, as an insect *Drosophila* is similar enough to humans to allow for translational research and at the same time sufficiently distinct from humans and other mammals to allow for the development of selective pest and disease vector control strategies that take advantage of their specific appetites. Furthermore, the full metamorphosis of *Drosophila* offers both its larval and adult stages, respectively dominated by feeding versus sexual motivation, as study cases for starvation-induced changes.

The effects of starvation in *Drosophila* are studied at many levels (Mirth and Riddiford, 2007; Partridge et al., 2005; Rion and Kawecki, 2007; Sakagiannis et al., 2025; Suarez-Grimalt et al., 2024; for larger flies: Dethier, 1976). As with behavioural adaptations upon starvation, the biogenic amines have turned out to be important mediators of nutritional-state information (Selcho and Pauls, 2019; Suarez-Grimalt et al., 2024). Of particular relevance within the central brain are the aminergic innervations of the mushroom bodies, prominent paired brain structures in arthropods, comprising intrinsic neurons called Kenyon cells (KCs) with characteristically elongated axons arranged as densely packed, parallel fibres. The mushroom bodies serve as a hub to integrate i) a sparse, combinatorial representation of the sensory environment including odours (provided to the KC dendrites by sensory projection neurons: PNs), ii) a representation of the ’matters of concern’ including food (provided by aminergic neurons intersecting the KC axon fibre bundle), and iii) a network of mushroom body output neurons (MBONs) that, via multiple synaptic steps, shape ongoing behaviour (**Fig. S1**). The mushroom bodies are involved in many forms of adaptive behaviour (Suarez-Grimalt et al., 2024), but in the present context it is particularly relevant that when odour-induced activity in a KC coincides with a dopaminergic reward signal elicited by for example sugar, presynaptic plasticity at the KC::MBON synapses is induced that is the basis for learned attraction when the odour is encountered again (adult flies: Davis, 2023; Li et al., 2020; Modi et al., 2020; larvae: Eschbach and Zlatic, 2020; Thum and Gerber, 2019). It is such odour-sugar reward associations that are typically used to investigate how starvation affects memory in *Drosophila*.

Indeed, when introducing the odour-sugar association paradigm for adult flies, Tempel et al. (1983) already reported that odour-sugar association leads to olfactory short-term memory in starved but not in sated animals, using an 18-22-h starvation protocol in culture vials equipped only with a wet filter paper (’wet starvation’). They also reported that neither odour preference in experimentally naïve animals nor memory after odour-shock training was affected by starvation, which is striking given that even at only slightly longer starvation periods the fitness and survival of the flies are compromised. Regardless, the requirement of starvation for robust odour-sugar association but not for odour-shock association has been much-replicated in the decades since this initial report (e.g. Ichinose and Tanimoto, 2016; Coban et al., 2024). Indeed, this work has been extended by Liu et al. (2012), who showed that starvation is required for associations of odour with the induced activation of a relatively broad set of rewarding dopaminergic/ serotonergic neurons (defined by the DDC-Gal4 driver strain), whereas it is not required for associations of odour with a likewise relatively broad set of punishing dopaminergic neurons (defined by TH-Gal4). However, it was shown by Gruber et al. (2013) and more recently by Meschi et al. (2024) that the robust odour-shock associations that can be observed even in sated flies can still be improved by starvation. In other words, according to these reports, starvation improves short-term memory for both odour-sugar *<u>and</u>*for odour-shock associations. This challenges what we here call the ’adaptive specificity hypothesis’, the notion that starvation selectively enhances food-related processing (Dethier, 1976; Sakagiannis et al., 2025; Selcho and Pauls, 2019; Sgammeglia and Sprecher, 2022; Suarez-Grimalt et al., 2024).

In this context, we survey a range of different association tasks for their starvation-sensitivity. In accordance with the studies referenced above, these tasks all address associative olfactory short-term memory and its modulation by 18-22 h wet starvation, but differ in terms of i) the reinforcers and ii) the number of training trials that were used, iii) the life stage of the experimental animals, iv) the predictive structure and associative timing of the task, v) whether memory is expressed as an increase or decrease in odour preference, and vi) whether the learned behaviour is motivated by the pursuit of reward or by the avoidance of/ escape from punishment. Whereas our results are in line with the adaptive specificity hypothesis with respect to the distinction between odour-sugar and odour-shock associations, we uncover additional complexity in starvation-sensitivity across the 26 tasks employed.

## Materials and Methods

The present study probes for the effects of starvation on associative short-term memory in adult and larval *Drosophila*, using 26 kinds of differential conditioning tasks for odours as conditioned stimuli. We did not attempt to ’streamline’ procedures across these tasks to achieve an arguably elusive uniformity. Rather, we adopted procedures corresponding to previously published studies from our own and other labs using these tasks, and opted for consistency within the present study only when this did not compromise the interpretation of our results in relation to these earlier studies.

### Fly husbandry and genotypes

*Drosophila melanogaster* were reared as mass cultures in vials of 4.8 cm diameter with standard cornmeal-molasses food at 60%-70% relative humidity, at 25 °C temperature, and under a 12 h:12 h light: dark cycle, unless mentioned otherwise. Behavioural experiments were conducted with mixed-sex cohorts either of flies aged 2-5 days after adult hatching, or of larvae aged 4 or 5 days after egg laying as mentioned in the *Results*. In the case of adult flies, scores were taken separately for females and males; as we did not observe significant differences in behaviour between the sexes (**Fig. S2-S7**), their data were combined throughout. In the case of the larvae, no attempt was made to determine their future sex, as somatically these animals are not yet sexually differentiated.

The Canton-S strain was used as wild-type. For optogenetic activation experiments, the experimental genotypes were generated by crossing a GAL4 driver strain covering the PPL1-01 neurons (PPL1-γ1pedc) (MB320C-Gal4, BDSC 68253; Aso et al., 2014) to the UAS-ChR2-XXL effector strain (UAS-ChR2-XXL, BDSC 58374; Dawydow et al., 2014). This yielded as offspring flies of the experimental genotype double heterozygous for the driver and the effector construct (in the following denoted as PPL1-01>ChR2XXL), allowing for optogenetic activation of the PPL1-01 neurons. Culture vials with experimental genotypes were shielded with black cardboard to avoid optogenetic activation by room light.

### Experiments with adult flies

#### Sated versus starved flies

Approximately 100 flies were randomly selected as mixed-sex cohorts from their culture vials and transferred either to fresh culture vials (Sated) or to vials without food but only a tissue paper soaked with tap water (Starved). After 18-22 h, the flies were collected from these vials for behavioural experiments (Fig. 1A).

**Fig. 1.**
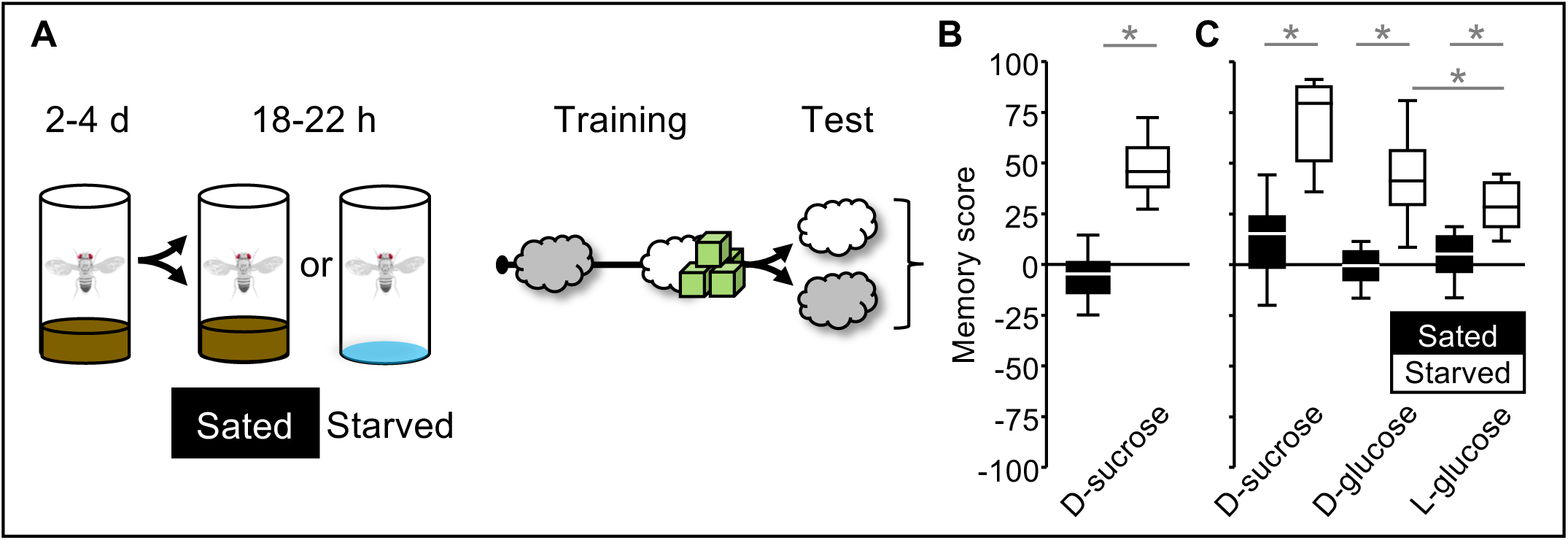
Starvation improves odour-sugar association. **(A)** Sated or starved wild-type flies were differentially conditioned in cohorts of ∼100 flies such that during training, one of two odours (white and grey clouds) was presented with a sugar reward (green cubes), followed by a choice to determine the relative preference for both odours by counting of the number of flies in either arm of the test T-maze. Based on these numbers, associative-memory scores were calculated according to Eqn 1 and Eqn 2, with positive and negative scores indicating relative preference for and avoidance of the trained odour, respectively. The chemical identity of the odours was balanced across repetitions of the experiment. **(B)** Relative to sated flies (filled boxplot), memory scores for 2 M of D-sucrose as the reward were increased in starved flies (open boxplot) (MWU test: U=4, p<0.05, N=24, 24). **(C)** A repetition of the experiment in (B) for three different sugars showed that relative to sated flies memory scores were increased in starved flies (D-sucrose, MWU tests: U=13, p<0.05, N=20, 20; D-glucose, MWU tests: U=28, p<0.05, N=20, 20; L-glucose, MWU tests: U=34, p<0.05, N=20, 20; starved D-glucose vs. L-glucose MWU tests: U=120, p<0.05, N=20, 20). * and ’ns’ respectively indicate significance and non-significance in MWU tests where α was adjusted with a Bonferroni–Holm correction to keep the experiment-wide type I error rate at <0.05. Statistical results, including OSS tests of memory scores against chance levels (i.e. zero), are reported along with the source data in the supplemental data file “Starvation DATA”. For the underlying preference scores and memory scores separated by sex, see **Fig. S2**.

#### Association of odour and sugar reward

We used an experimental set-up as initially described by Tully and Quinn (1985), modified as in Schwaerzel et al. (2003) (CON-ELEKTRONIK, Greussenheim, Germany), at 20-25°C and 60-80% relative humidity. In brief, flies were differentially conditioned such that two odours were presented to the flies during training, but only one of these odours was paired with a sugar reward (+). This was followed by a choice test between the two odours presented from the two arms of a T-maze. Training was performed under white light, testing in red light that is invisible to the flies.

Undiluted 50 μl benzaldehyde (BA) and 250 μl 3-octanol (3OCT) (CAS 100-52-7 and 589-98-0, respectively; Merck, Darmstadt, Germany) were used as odorants, applied to 1-cm-deep Teflon containers of 5 mm and 14 mm diameter, respectively. The use of BA and 3OCT as odorants was balanced across repetitions of the experiment, as was the sequence of trial types during training (when BA was rewarded: 3OCT/BA+ or BA+/3OCT, and for the reciprocal cases when 3OCT was rewarded: BA/3OCT+ or 3OCT+/BA).

D-sucrose, D-glucose and L-glucose were used as a sugar reward as described in Yarali and Gerber (2010) and König and Gerber (2022). Specifically, for rewarded trials, the training tube was lined with filter paper that had been soaked the day before with 1.25 ml of 2 mol/l of these sugars (D-sucrose CAS: 57-50-1, Roth, Karlsruhe, Germany; D-glucose CAS: 50-99-7; L-glucose CAS: 921-60-8), as indicated in the *Results*, and left to dry overnight. The same was done for non-rewarded trials, except that sugar was omitted.

Flies were loaded to the choice point and after 1 min released into a training tube without reward (or a training tube with reward), while one of the odours was simultaneously presented for 45 s. Then the flies were moved back to the choice point and 1 min later released into a training tube with the other reward condition, and the other odour was presented for 45 s, after which the flies were moved to the choice point again. After approximately 3 min, the flies were allowed to choose between the two odours presented on either side from the choice point. After 2 min, the arms from the choice point were closed, and the flies in each arm of the T-maze were counted. From these numbers, the preference score was calculated as:

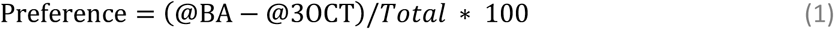

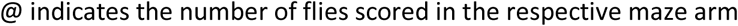

Preference scores thus range between 100 and -100, with positive values indicating a relative preference for BA and negative values indicating a relative preference for 3OCT.

As mentioned, pairing of BA and 3OCT with the sugar reward (BA+ and 3OCT+, respectively) was balanced across repetitions of the experiment. This allows an associative memory score to be calculated as the difference in preference between these reciprocally trained sets of flies:

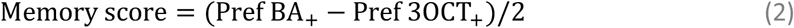

Thus, memory scores also range between 100 and -100, with positive values indicating a relative preference for the sugar-associated odour while negative values would indicate its avoidance.

### Six-trial association of odour and electric shock

Six-trial association experiments of odour with electric shock were in principle performed as described in the preceding section, with modifications as mentioned below and corresponding to previously published work (Yarali et al., 2008). Electric shocks were applied via copper grids inside the training tubes as a 2-min train of 24 shock pulses of 100 V, with each pulse lasting for 1.2 s and followed by a 3.8 s off-period. Flies were loaded into the training tubes and after 1 min one of the odours was presented for 15 s, in all cases. In the punishment-learning version of the paradigm, at 5:00 min the other odour was presented for 15 s, and upon odour offset electric shock was delivered at 5:15. This temporal relationship corresponds to an inter-stimulus interval (ISI) of -15 s, defined as the time point of odour onset minus the time point of shock onset. For the relief-learning version of the paradigm, the odour was presented at an ISI of +145 s. In other words, for punishment learning odour onset precedes shock onset by 15 s (negative ISI), whereas for relief learning shock onset precedes odour onset by 145 s (positive ISIs), thus leaving a 25 s gap between shock offset and odour onset. At 10:00 min, the flies were removed from the set-up into vials corresponding to their treatment condition (Sated: vials with food, Starved: vials with wet tissue paper) and kept there for 16 min until the next of a total of six training trials ensued. For testing, the flies were loaded to the choice point, and testing and calculations of preferences and memory scores were carried out as described in the preceding section.

### One-trial association of odour and electric shock

One-trial association experiments of odour with electric shock were in principle performed as described in the preceding section, with the modifications mentioned below and corresponding to previously published work (Galili et al., 2011; Shuai et al., 2011; Tully and Quinn, 1985). Flies were loaded into the training tubes and after 1 min one of the odours was presented for 15 s. The other odour was then presented for 15 s at one of six ISIs (-75, -60, -45, -30, -15, or 0 s) relative to electric shock, which was delivered from 4:45 min on as a 1-min train of 12 shock pulses. Given these temporal relationships between odour and shock, the -75 to -30-s ISIs are defined as ’trace’, the -15-s ISI as ’delay’, and the 0-s ISI as ’simultaneous’ conditioning (Galili et al., 2011). At 7:00 min, the flies were transferred to the choice point, and testing and calculations of preferences and memory scores were carried out as described above. The sequence of trial types during training was as described in half of the cases; in the other half of the cases it was the reverse, such that the shock-associated odour, at the indicated ISIs, was presented first, with shock onset at 1:45 and presentation of the other odour from 4:00 min on.

### Association of odour and PPL1-01 activation

Experiments on the association of odour with optogenetic activation of the PPL1-01 dopaminergic neurons were in principle performed as described in the preceding section, with the modifications mentioned below and corresponding to previously published work (Amin et al., 2025). Training tubes were used equipped with 24 LEDs emitting light at a peak wavelength of 465 nm (blue light) mounted on the outer surface of the training tubes. Blue light stimulation involved 1-min trains of 12 pulses with each pulse lasting for 1.2 s, followed by a 3.8 s off-period. The absolute irradiance at the centre of the training tube during stimulation was 200 μW/cm^2^, as measured with an STS-VIS spectrometer (Ocean Optics, Ostfildern, Germany).

Flies of the PPL1-01>ChR2XXL genotype, allowing for optogenetic activation of the PPL1-01 neurons, were loaded to the training tubes, and after 1 min one of the odours was presented for 1 min, in all cases. The other odour was presented for 1 min at one of four ISIs (-100, -15, 120, or 240 s) relative to light stimulation, which was delivered from 6:35 on. Given these temporal relationships between odour and optogenetic activation, the -100-s ISI is regarded as ’trace’, the -15-s ISI as ’delay’, and the further ISIs as ’relief’ conditioning. At 15:00 min, the flies were transferred to the choice point, and testing and calculations of preferences and memory scores were carried out as mentioned above.

We additionally performed an experiment with modified procedures to accommodate six-trial relief conditioning with PPL1-01 activation. In this case, light stimulation was delivered from 4:15 on, and an ISI of 120 s was used. At 9:00 min, the flies were removed from the set-up into vials corresponding to their treatment condition (Sated: vials with food, Starved: vials with wet tissue paper) and kept there for 16 min until the next of a total of six training trials ensued. For testing, the flies were loaded to the choice point, and after 5 min testing and calculations of preferences and memory scores were carried out as described above.

### Extinction of an odour-shock association

Protocols to measure extinction memory were adapted from Felsenberg et al. (2018) and Schwaerzel et al. (2003) and were performed in principle as described in the section *One-trial association of odour and electric shock* with the modifications described below. After single-trial differential conditioning with odours and electric shock, the flies were removed from the set-up into vials corresponding to their treatment condition (Sated: vials with food, Starved: vials with wet tissue paper). After 30 min, the flies were loaded into transparent tubes without a copper grid that were positioned within the set-up as in training. After 3 min, the flies in the baseline condition were exposed to a blank stimulation, i.e. empty odour cups were fitted to the set-up for 15 s, twice with a 15 min gap, whereas for extinction training, the flies were exposed to the previously shock-associated odour on these occasions. Once a total of 70.5 min had passed after the training trial, the flies were transferred to the choice point, followed by testing and the calculation of preferences and memory scores as described above. In addition, we quantified extinction memory as follows:

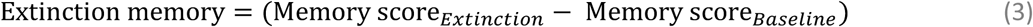

Thus, positive extinction memory scores indicate that extinction learning of the odour-shock association had taken place; negative extinction memory scores would indicate a paradoxically enhancing effect on the odour-shock association if the odour is presented alone.

### Experiments with larvae

#### Sated versus starved larvae

Larvae were starved for a period either from 4 days after egg laying to 5 days, or one day earlier (from 3 to 4 days), such that starvation was induced respectively either after or before the larvae were likely to have passed the critical weight for pupariation (Stieper et al., 2008; Mirth et al., 2005). Specifically, following a protocol from Brünner et al. (2020), approximately 60 larvae aged 4 days (or 3 days) after egg laying and of either prospective sex were randomly selected from their culture vials and transferred to 2.8-cm-diameter vials filled with solidified 2.5% agarose to a depth of 1 cm (electrophoresis grade; CAS: 9012-36-6, Roth), to which a water droplet of approximately 40 µl was added to prevent the dehydration of the larvae. After approximately 20 h, the larvae that were now 5 days old (or 4 days old) (Starved) were collected for behavioural experiments and compared to non-starved larvae of the same age (Sated).

#### Association of odour and sugar reward

Experiments on the association of odour and sugar reward (+) used a three-trial differential conditioning paradigm that was modified to measure the effects of paired and unpaired presentations of odour and sugar separately (Schleyer et al., 2018). As odorants, n-amyl acetate (AM) (CAS: 628-63-7, Merck) diluted 1:20 in paraffin oil (CAS: 8042-47-5, AppliChem), and undiluted 1-octanol (1OCT) (CAS: 111-87-5; Merck) were used. These were presented in custom-made Teflon containers from which odour could evaporate. As a sugar reward (+), 2 mol/L D-fructose (CAS: 57-48-7; Roth) was used in 1% agarose.

Approximately 20 larvae were placed in the middle of a Petri dish (85 mm in diameter for the 5-day-old and 55 mm for the smaller, 4-day-old larvae) containing 1% solidified agarose and 1OCT as the odour, presented from two odour containers located on opposite sides of the Petri dish (1OCT). After 2.5 min, the larvae were transferred to a fresh Petri dish containing agarose with the sugar reward added, and odour containers with AM, and left there for another 2.5 min (AM+). Three such cycles of training with 1OCT as the unpaired odour and AM as the paired odour were performed, in each case using fresh Petri dishes. In half of the cases, training started with 1OCT followed by AM+ (1OCT/ AM+), whereas in the other half of the cases, the sequence was the reverse (AM+/1OCT). Following training, the larvae were transferred to the middle of a test Petri dish and tested for their preference for AM by placing a container with AM on one side of the Petri dish and an empty (EM) container on the other side. After 3 min, the number of larvae on either side was counted and the preference score was calculated as:

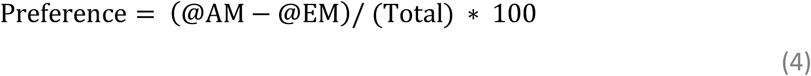

Positive values indicate a preference for AM, and negative values an avoidance of AM. For each cohort of larvae for which AM was the paired odour during training, a second cohort was trained with reciprocal contingencies, such that AM was the unpaired odour and 1OCT the paired odour (AM/1OCT+, or 1OCT+/AM), followed by testing for AM preference as detailed above. Thus, AM preference scores were taken after AM had been presented during training paired with sugar or unpaired from it.

Interpreting the AM preferences that result from paired versus unpaired training requires a measure of baseline AM preference cleared of associative effects. For larval *Drosophila* this can be achieved by an experimental twist exploiting the nature of learned appetitive behaviour as a search, which adaptively ceases in the presence of the sought-for item (Gerber and Hendel, 2006): when larvae are trained as detailed above, but the test is carried out in the presence of the sugar reward, the larvae show the same level of AM preference regardless of whether AM has been trained in a paired or in an unpaired manner (Schleyer et al., 2018). In other words, under such test conditions AM preferences are cleared of the associative effects of training (’innate’ AM preferences are not affected by the presence of sugar: Schleyer et al., 2011). Thus, the larvae were differentially conditioned as detailed above such that AM was either the paired odour or the unpaired odour, followed by a test for their AM preference; however, to assess baseline AM preferences the test was performed, in independent sets of larvae, in the presence of sugar. Whenever, as previously reported (Saumweber et al., 2011; Schleyer et al., 2018; Sen et al., 2024), the preference values did not differ under the corresponding test conditions, the data were pooled and their median value was used as the Baseline. This allowed a Paired and Unpaired memory score to be calculated as:

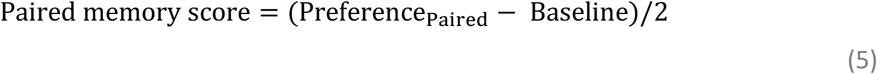

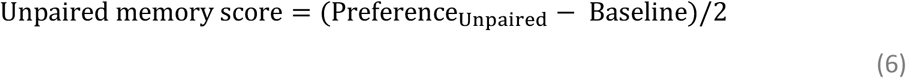

Thus, positive scores indicate a higher-than-baseline preference for the odour, whereas negative scores indicate a lower-than-baseline preference for the odour. These types of memory reflect the fact that the animals have learned about AM as a predictor of the presence versus the absence of the reward, respectively (Sen et al., 2024) (in psychological terminology ’conditioned excitation’ versus ’conditioned inhibition’: Rescorla, 1988).

### Association of odour and bitter-taste punishment

The experiments were in principle performed as described in the preceding section, except that instead of the sugar reward, 5 mM quinine (CAS: 207671-44-1; Sigma-Aldrich) in 1% agarose was used as a taste punishment. Notably, learned aversive behaviour can be characterized as an attempt to escape, which adaptively is not performed in the absence of the taste punishment (Gerber and Hendel, 2006; Schleyer et al., 2011; Paisios et al., 2017). In other words, associations of odour with taste punishment are behaviourally expressed in the presence, but not in the absence of the taste punishment (’innate’ AM preferences are not affected by the presence of quinine: Schleyer et al., 2011).

### ’Innate’ preference tests for adult flies and larvae

#### Sugar preference in adult flies

Sated or starved flies were collected from their vials for an immediate test of their sugar preference (Fig. 10A). 2M D-sucrose, D-glucose or L-glucose was used as a sugar, presented on dried filter paper as described in the section *Association of odour and sugar reward*. The flies were loaded to the choice point of the T-maze and allowed to choose between a sugar side or a blank arm of the maze. After 2 min the arms of the maze were closed, and the number of flies in each arm was counted to calculate the preference score according to Equation 4, with due modification. Positive values indicate a preference for, and negative values an avoidance of sugar.

### Odour preference in larvae

Sated or starved larvae of the indicated age were collected from their vials for an immediate test of their odour preference. As odorants, either n-amyl acetate (AM) (Fig. 10B, Fig. 10J) or 1-octanol (1OCT) (Fig. 10C, Fig. 10K) was used, and odour preference was determined in a Petri dish assay as described for Equation 4, with due modification, such that positive values indicate a preference for, and negative values an avoidance of the respective odour. Petri dishes of 85 mm diameter were used for the 5-day-old larvae, whereas 55-mm-diameter Petri dishes were used for the smaller, 4-day-old larvae.

### Fructose and quinine preference in larvae

Sated or starved larvae of the indicated age were collected from their vials for an immediate test of their preference for either fructose or quinine. Approximately 20 larvae were placed in the middle of a Petri dish containing only 1% solidified agarose on one side and D-fructose or quinine in 1% agarose on the other side. After 3 min, the number of larvae on either side was counted, and the preference score was calculated according to Equation 4, with due modification, such that positive values indicate a preference for, and negative values an avoidance of the respective tastant. Petri dishes of 85 mm diameter were used for the 5-day-old larvae, whereas 55-mm-diameter Petri dishes were used for the smaller, 4-day-old larvae.

### Locomotion analysis

Animals were placed in groups of 7-8 in the centre of Petri dishes (85 mm inner diameter), filled with 1% agarose. The Petri dishes were placed in a ring with infrared LEDs for illumination (850 nm; Solarox), and the behaviour of the animals was videorecorded for three minutes with a camera (Basler acA2040-90um). Afterwards, the behaviour was analysed using IMBA (Individual Maggot Behaviour Analysis) as described in Thane et al. (2023). This system can track the identity of individual larvae across collisions and is based on the definition of 12 equidistant spine points from the head to the tail of each larva, as well as vectors between specific spine points to characterize the front (head vector) and rear (tail vector) of the animal.

As it has been shown previously that the crawling speed of larvae depends directly on their body size (Aleman-Meza et al., 2015; Thane et al., 2023), we normalized all measurements of distance or velocity to the individual body length (bl) of each animal. The body length was calculated as the length of the spine of an animal, averaged over the complete observation period.

To visualize the animals’ behaviour, all tracks corresponding to a given condition were first plotted on top of each other (Fig. 10F, Fig. 10G, Fig. 10N, Fig. 10O); then five non-overlapping sample tracks of at least 90 s duration were randomly selected from each condition (Fig. 10H, Fig. 10I, Fig. 10P, Fig. 10Q).

The occurrence of strong lateral head movements, so-called head casts (HC), is indicated in red and green for right and left HCs, respectively. The definition of an HC is based on the occurrence of an absolute head vector angular speed (see below) above 35 °/s. For details, see Thane et al. (2023).

We further analysed the behaviour of all individual animals that were tracked for at least half of the total video time (90 s). This ensures that each individual can enter the analysis only once. In rare cases, if the head and tail of an animal could not be correctly assigned, the animal was excluded from the analysis; this was the case for 21 out of 509 animals. For each animal included in the analysis, we determined the following behavioural attributes:

- Head velocity (bl/s): the mean velocity of the head point in a forward direction (that is, in the direction in which the body is oriented) over the complete observation period. Thus, if a larva moves relatively straight and does little ’zigzagging’, values for this attribute are high.
- Absolute head vector angular speed (°/s): the mean absolute angular speed of a vector through the front part of the animal (the head vector) over the complete observation period, indicating how much the head moves sideways, independent of direction. Thus, if a larva moves relatively straight and does little ’zigzagging’, values for this attribute are low.

As a larva can increase its speed during a relatively straight ’run’ either by increasing the Inter-step distance, i.e. by taking larger steps, and/or by decreasing the Inter-step interval, i.e. by taking less time for each step, we further determined:

- Inter-step distance (bl): the mean distance that a larva covers between two ’steps’, that is, within each peristaltic cycle of forward movement.
- Inter-step interval (s): the mean time interval that a larva takes for each peristaltic cycle of forward movement.

### Statistical analyses

All the data were analysed using non-parametric statistics and are represented as boxplots, showing the median as the midline, the 25 and 75% quantiles as box boundaries, and the 10 and 90% quantiles as whiskers. To compare data across multiple independent experimental groups, Kruskal-Wallis (KW) tests were employed, followed by Mann-Whitney U (MWU) tests for subsequent pairwise comparisons. When assessing whether scores within individual experimental groups differed significantly from zero (i.e. chance levels), one-sample sign tests (OSS tests) were performed. To account for multiple tests conducted within a single experiment, a Bonferroni-Holm (Holm 1979) correction was applied to maintain an experiment-wide error rate of 5%. This involved dividing the critical p-value of <0.05 by the number of tests performed. Significant results are denoted by asterisks (*) for MWU tests. Correlation analyses used Pearson correlation coefficients. All statistical analyses were performed using Statistica (version 12; Statsoft Inc., Tulsa, OK, USA). Sample sizes were planned in accordance with prior, similar studies and are reported in the figure legends. The results of the statistical analyses and the source data of all experiments performed are documented in the supplemental data file ‘Starvation DATA’.

## Results

### Experiments with adult flies

#### Starvation improves odour-sugar association

Flies of either the sated or the starved state underwent one-trial differential conditioning, such that one odour was paired with 2 M of D-sucrose as the sugar reward, whereas the other odour was presented without the reward, followed by a choice between the two odours (**Fig. 1A**). Relative to sated flies, memory scores were higher in starved flies (**Fig. 1B**), with memory scores of sated flies remaining at chance levels (supplemental data file "Starvation DATA"). These results align with previous findings (e.g. Krashes et al., 2009; Tempel et al., 1983) and show that our starvation protocol effectively induced the expected and mnemonically relevant state change.

Our result for 2 M of D-sucrose was replicated and extended for 2 M of D-glucose as well as for 2 M of L-glucose (**Fig. 1C**), a sugar that is indistinguishable from D-glucose at the sensory level (in terms of its ’sweetness’), but which differs from D-glucose in that it is of no metabolic value to flies (Fujita and Tanimura, 2011). It is therefore of particular significance that in the starved state memory scores for 2 M of D-glucose are higher than for L-glucose (see *Discussion*).

### Starvation leaves six-trial ’punishment’ associations unaffected, but impairs ’relief’ associations

Sated or starved flies underwent differential conditioning and were compared for six-trial odour-shock ’punishment’ associative memory, and six-trial shock-odour ’relief’ associative memory (**Fig. 2A**). Relative to the sated condition, starvation did not impact aversive memory scores observed upon odour-shock punishment training (**Fig. 2B, left**). However, the appetitive memory scores observed upon shock-odour relief training were reduced, and indeed abolished, in starved flies (**Fig. 2B, right**). Thus, starvation has task-dependent effects on memory, in that memory may be improved (odour-sugar association, **Fig. 1**), impaired (shock-odour relief association, **Fig. 2B, right**), or unaffected (odour-shock punishment association, **Fig. 2B, left**) by starvation. Critically, therefore, neither the involvement of shock versus sugar, nor the question whether the memory scores in question are positive or negative, is the sole determinant for sensitivity to starvation. We therefore ventured to undertake a survey of the effects of starvation across 21 additional associative memory tasks, next focusing on one-trial odour-shock association.

**Fig. 2.**
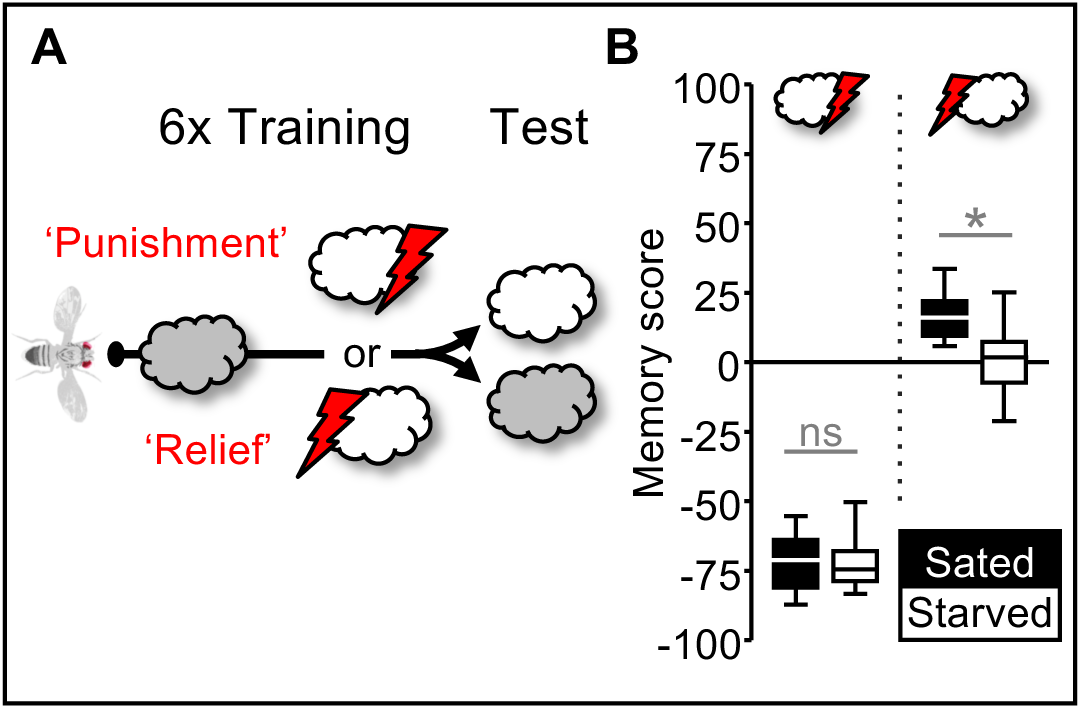
Starvation leaves six-trial ’punishment’ associations unaffected, but impairs ’relief’ associations. **(A)** Using a starvation protocol as in Fig. 1, cohorts of ∼100 sated or starved wild-type flies were differentially conditioned such that a reference odour (grey cloud) was presented, followed by the presentation of the trained odour (white cloud) either before (’Punishment’) or after (’Relief’) electric shock (red lightning bolt). After six such training trials, there followed a choice to determine the relative preference for both odours by counting the number of flies in either arm of the test T-maze. Based on these numbers, associative-memory scores were calculated according to Eqn 1 and Eqn 2, with positive and negative scores indicating relative preference for and avoidance of the trained odour, respectively. The chemical identity of the odours was balanced across repetitions of the experiment. **(B)** For odour-then-shock ’punishment’ conditioning, no difference in memory scores was found between sated and starved flies (filled and open boxplots, respectively) (MWU test: U=95, p=0.91, N=14, 14), whereas for shock-then-odour ’relief’ conditioning, memory scores were reduced in starved flies (MWU test: U=92, p<0.05, N=22, 22). * and ’ns’ respectively indicate significance and non-significance in MWU tests where α was adjusted with a Bonferroni–Holm correction to keep the experiment-wide type I error rate at <0.05. Statistical results, including OSS tests of memory scores against chance levels (i.e. zero), are reported along with the source data in the supplemental data file “Starvation DATA”. For the underlying preference scores and memory scores separated by sex, see **Fig. S3**.

### Starvation leaves one-trial odour-shock associations intact

We wondered whether an effect of starvation on odour-shock associations was obscured in the previous experiment by ceiling effects through overtraining in the six-trial regimen that we had used. To survey odour-shock associations across a range of memory strengths that would allow us to detect either increases or impairments of memory, we tested for the effect of starvation on one-trial odour-shock associations at different inter-stimulus-intervals (ISIs) between the onset of odour and the onset of shock (**Fig. 3A**). Specifically, we used ISIs of -75, -60, -45, -30s, -15 or 0 s. Given a 15-s odour presentation, for the ISIs between -75 and -30 s this leaves temporal gaps between odour offset and shock onset of 60, 45, 30, or 15 s, respectively (’trace’ conditioning), whereas for the -15 s ISI odour offset coincides with shock onset (’delay’ conditioning) and for the 0-s ISI odour onset coincides with shock onset (’simultaneous’ conditioning) (Galili et al., 2011; Shuai et al., 2011; Grover et al., 2022). As expected, memory scores were stronger the closer in time odour and shock were presented, but critically in none of the cases did we observe an effect of starvation (**Fig. 3B**) (a trend at ISI -45 s could not be substantiated: **Fig. 3C**). This absence of an effect of starvation on trace conditioning, delay conditioning and simultaneous conditioning is not trivial, given that these forms of conditioning only partially overlap in their underlying mechanisms (Galili et al., 2011; Grover et al., 2022).

**Fig. 3.**
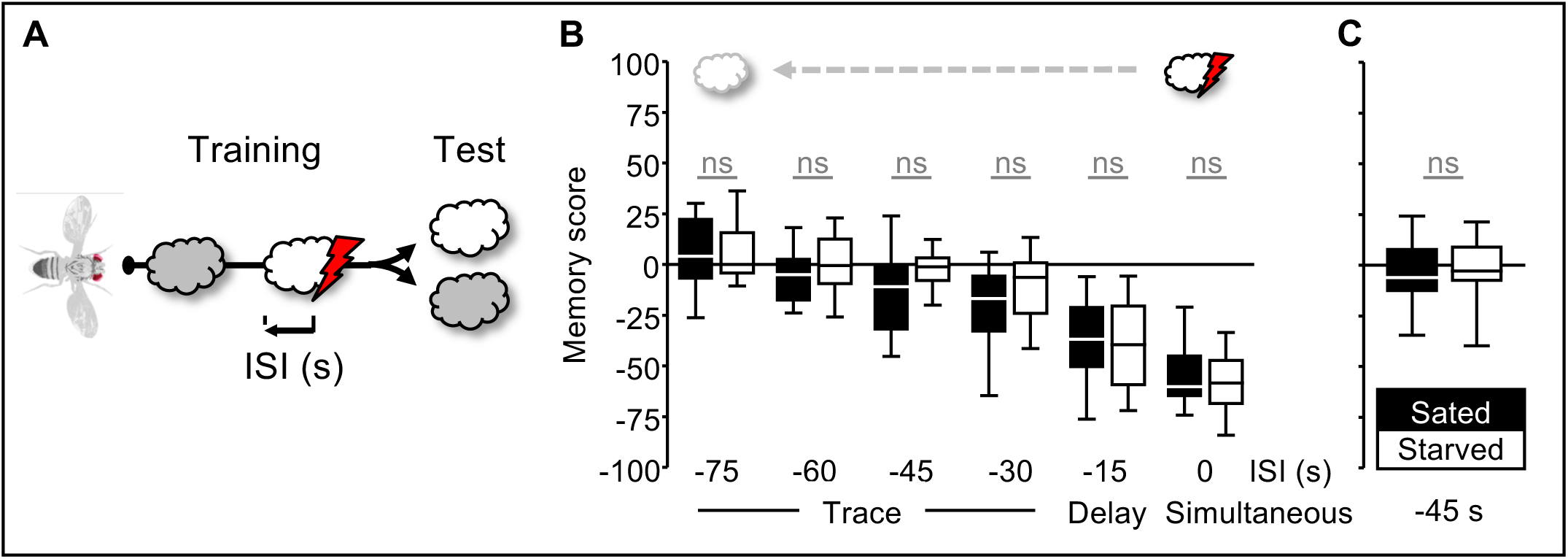
Starvation leaves one-trial odour-shock associations intact. **(A)** Using a starvation protocol as in Fig. 1, cohorts of ∼100 sated or starved wild-type flies were differentially conditioned such that a reference odour (grey cloud) was presented, followed by the presentation of the trained odour (white cloud) and electric shock (red lightning bolt) at one of the indicated inter-stimulus-intervals (ISIs), establishing trace conditioning (ISIs between -75s and -30s), delay conditioning (ISI=-15s) or simultaneous conditioning (ISI=0s). After one such training trial, there followed a choice to determine the relative preference for both odours by counting the number of flies in either arm of the test T-maze. Based on these numbers, associative-memory scores were calculated according to Eqn 1 and Eqn 2, with positive and negative scores indicating relative preference for and avoidance of the trained odour, respectively. The chemical identity of the odours was balanced across repetitions of the experiment. **(B)** Memory scores of sated and starved flies (filled and open boxplots, respectively) did not differ for any of the ISIs (MWU tests, from left to right: U=365, p=0.83, N=28, 27; U=355, p=0.32, N=28, 30; U=256, p=0.03, N=28, 28; U=297.5, p=0.18, N=27, 28; U=321, p=0.95, N=25, 26; U=270, p=0.30, N=25, 26). **(C)** A repetition of the experiment in (B) with ISI=-45s did not substantiate the trend for reduced memory scores in starved flies seen in (B) (MWU test: U=368, p=0.55, N=28, 29). ’ns’ indicates non-significance in MWU tests. For these MWU tests, α was adjusted with a Bonferroni–Holm correction to keep the experiment-wide type I error rate at p<0.05 across (B); in (C), p<0.05 was used without such a correction. Statistical results, including OSS tests of memory scores against chance levels (i.e. zero), are reported along with the source data in the supplemental data file “Starvation DATA”. For the underlying preference scores and memory scores separated by sex, see **Fig. S4**.

### Starvation leaves odour-PPL1-01 associations intact

Associations of odours and electric shock in *Drosophila* involve the coincidence of olfactory processing and shock-evoked dopamine signals in the mushroom body (Li et al., 2020; Modi et al., 2020; Davis, 2023) (**Fig. 4A**). While electric shock activates multiple types of dopaminergic mushroom body input neurons (DANs; Riemensperger et al., 2005; Mao and Davis, 2009; Dylla et al., 2017), single, properly timed presentations of odours with the opto- or thermogenetic activation of the PPL1-01 type of DANs can establish trace, delay and relief associations (Tanaka et al., 2008; Aso et al., 2010; Hige et al., 2015; König et al., 2018; Aso et al., 2019; Aso et al., 2012). In other words, although the temporal ’fingerprint’ of association formation with different ISIs is qualitatively similar for electric shock and PPL1-01 activation, PPL1-01 provides but a subset of the dopaminergic signals triggered by shock. We were therefore curious to see whether trace, delay or relief memories established by PPL1-01 activation have starvation-sensitive components. To this end, sated or starved flies underwent one-trial training with odour and PPL1-01 activation at inter-stimulus-intervals (ISIs) establishing trace (-100 s), delay (-15 s) or relief associations (120, 240 s) (**Fig. 4B**), leaving a 40-s temporal gap between odour offset and PPL1-01 activation for the trace ISI, no gap for the delay ISI, and a 60-s or a 180-s gap for the two relief ISIs. Critically, in none of these cases did we observe an effect of starvation (**Fig. 4C**). Given that in our earlier experiment (**Fig. 2B, right**), an impairment in shock-relief associations was observed with six training trials, we repeated the experiment for PPL1-01 relief, this time likewise using six training trials, but did not observe an impairment by starvation either (**Fig. 4D**) (indeed, if any trend were to be gleaned from these data, it would be that PPL1-01 relief associations were increased, rather than impaired, in starved flies). Thus, relief associations established by shock are susceptible to impairment by starvation (**Fig. 2B, right**), but relief associations established by PPL1-01 are not (**Fig. 4C**, **Fig. 4D**).

**Fig. 4.**
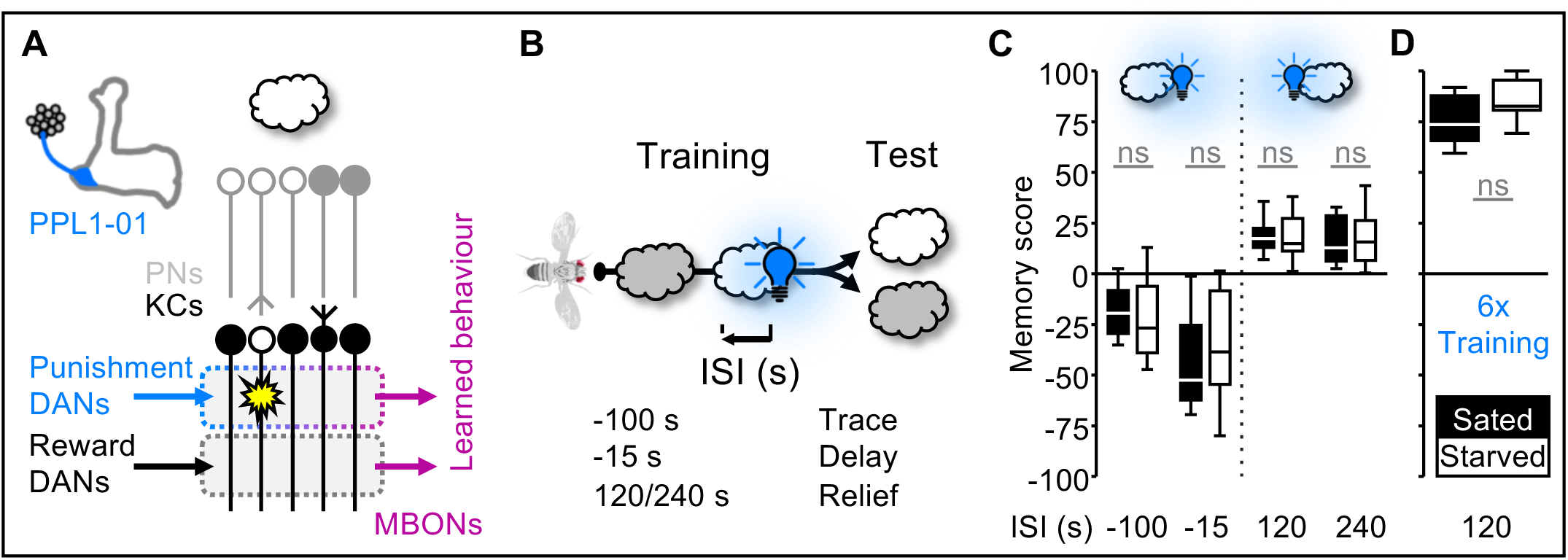
Starvation leaves odour-PPL1-01 associations intact. **(A)** Schematic morphology of the PPL1-01 dopaminergic neurons (DANs) (modified from Aso and Rubin, 2016) and simplified sketch of the circuitry underlying odour-DAN association (for references see body text). The combined divergence-convergence connectivity of olfactory projection neurons (PNs) and mushroom body intrinsic neurons (Kenyon cells, KCs) establishes a sparsened, combinatorial representation of the odour (open cloud, PN and KC cell bodies). Punishing dopaminergic DANs such as PPL1-01 intersect the axons of the mushroom body neurons within confined compartments (stippled box, top). Associative coincidence of odour activation and signalling from PPL1-01 induces presynaptic plasticity (star) at the synapses from odour-activated KCs to the compartmental mushroom body output neuron (top MBON). No such plasticity takes place in compartments receiving input from rewarding DANs (bottom MBON). Upon encountering the odour again, the changed balance of activity across the MBONs leads to net avoidance as the learned behaviour. **(B)** Using a starvation protocol as in Fig. 1, cohorts of ∼100 sated or starved flies were differentially conditioned such that a reference odour (grey cloud) was presented, followed by the presentation of the trained odour (white cloud) and blue light for optogenetic PPL1-01 activation (blue light bulb) at one of the indicated inter-stimulus-intervals (ISIs), establishing trace conditioning (ISI=-100s), delay conditioning (ISI=-15s) or relief conditioning (ISIs=120 or 240s). After one such training trial, there followed a choice to determine the relative preference for both odours by counting the number of flies in either arm of the test T-maze. Based on these numbers, associative-memory scores were calculated according to Eqn 1 and Eqn 2, with positive and negative scores indicating relative preference for and avoidance of the trained odour, respectively. The chemical identity of the odours was balanced across repetitions of the experiment. **(C)** Memory scores of sated and starved flies (filled and open boxplots, respectively) did not differ for any of the ISIs (MWU tests, from left to right: U=164, p=0.34, N=20, 20; U=138, p=0.22, N=19,19; U=215, p=0.72, N=23, 20; U=152, p=0.99, N=17, 18). **(D)** A modified repetition of the experiment in (C), but this time with six trials of relief conditioning (ISI=120s), did not uncover a difference in memory scores between sated and starved flies either (MWU test: U=75, p=0.31, N=14,14). ’ns’ indicates non-significance in MWU tests. For these MWU tests, α was adjusted with a Bonferroni–Holm correction to keep the experiment-wide type I error rate at p<0.05 across (C); in (D), p<0.05 was used without such a correction. Statistical results, including OSS tests of memory scores against chance levels (i.e. zero), are reported along with the source data in the supplemental data file “Starvation DATA”. For the underlying preference scores and memory scores separated by sex, see **Fig. S5**.

### Starvation leaves extinction of an odour-shock association intact

An important feature of memory systems is that after memories are formed there remains flexibility in the light of further experience. Specifically, if after a training phase during which an odour-shock association is established the odour is presented alone, flies do not erase the initially acquired odour-shock association but rather establish what is called an ’extinction memory’ for the odour, such that odour avoidance does not continue once the flies realize that the odour-shock contingency is broken (Felsenberg et al., 2018; Schwaerzel et al., 2003). Given that the initial acquisition of odour-shock memory is intact upon starvation, we wondered in turn whether extinction memory is affected. Sated or starved flies underwent one-trial differential conditioning followed either by two blank presentations of only air (Baseline) or by two presentations of the previously punished odour (Extinction), allowing an assessment of extinction memory as the difference in performance between these two conditions (**Fig. 5A**, **Fig. 5B**). Relative to the sated condition, starvation impacted neither memory scores in the baseline condition (**Fig. 5B**) nor extinction memory (**Fig. 5C**). We note that according to our experimental protocol memory scores in the baseline condition reflect an approximately 1-h memory.

**Fig. 5.**
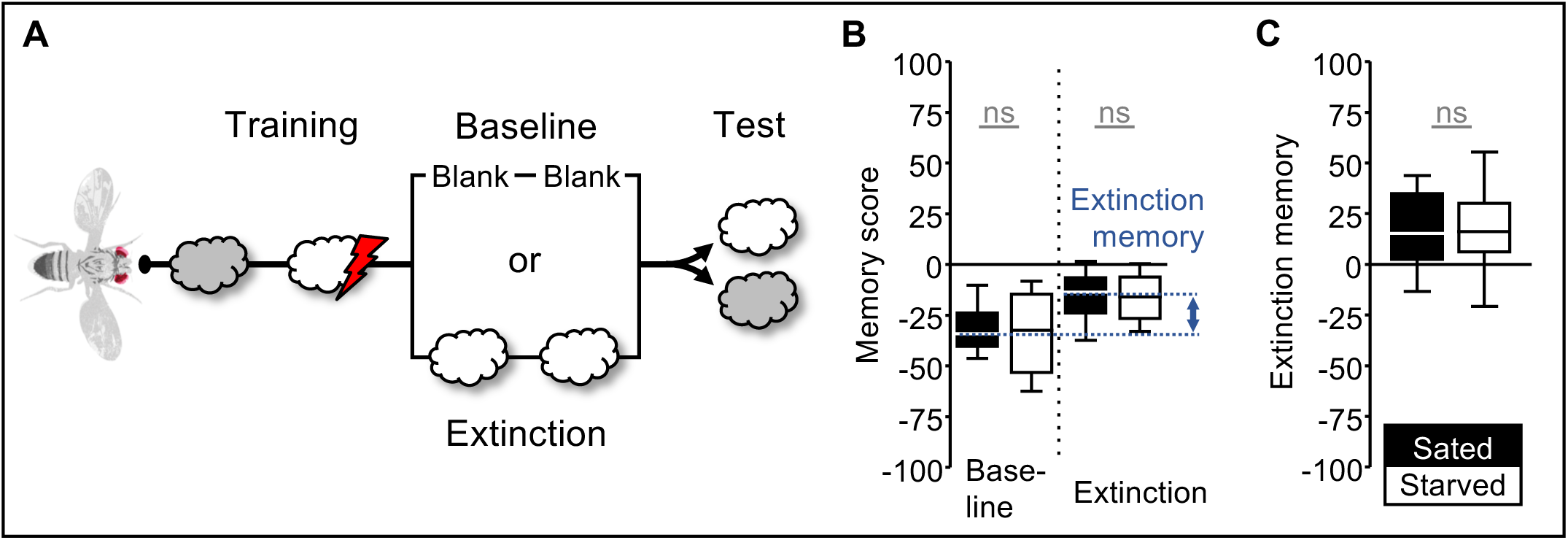
Starvation leaves extinction of an odour-shock association intact. **(A)** Using a starvation protocol as in Fig. 1, cohorts of ∼100 sated or starved wild-type flies were differentially conditioned such that a reference odour (grey cloud) was presented, followed by the presentation of the trained odour (white cloud) and electric shock (red lightning bolt) at an inter-stimulus interval of -15 s. After one conditioning trial, the flies were removed from the setup into vials corresponding to their treatment condition (Sated: vials with food, Starved: vials with wet tissue paper). After 30 min, the flies were returned to the setup and exposed either to two blank trials separated by 15 min (Baseline), or to the previously shock-associated odour (Extinction), followed by a choice test between the two odours. Based on the number of flies on each side, associative-memory scores were calculated according to Eqn 1 and Eqn 2, with positive and negative scores indicating relative preference for and avoidance of the shock-associated odour, respectively. Extinction memory is calculated according to Eqn 3. **(B)** No difference in memory scores was found between sated and starved flies (filled and open boxplots, respectively), either in the baseline condition (MWU test: U=48, p=0.91, N=10, 10) or in the extinction condition (MWU test: U=49, p=0.97, N= 10, 10). **(C)** Extinction memory did not differ between sated and starved flies (MWU test: U=48, p=0.91, N=10, 10). * and ’ns’ respectively indicate significance and non-significance in MWU tests where α was adjusted with a Bonferroni–Holm correction to keep the experiment-wide type I error rate at <0.05. Statistical results, including OSS tests of memory scores against chance levels (i.e. zero), are reported along with the source data in the supplemental data file “Starvation DATA”. For the underlying preference scores and memory scores separated by sex, see **Fig. S6**.

As a summary of the results in adult flies, we plotted the median memory scores of sated versus starved flies (**Fig. 6A**), revealing odour-sugar associations and shock-relief associations as the apparently exceptional cases among the 18 tasks studied. Interestingly, although both these associations support positive memory scores, the effects of starvation are opposite in the two cases: starvation improved odour-sugar associations but impaired shock-relief associations. We further note that median memory scores in females and males were virtually identical throughout (**Fig. 6B**).

**Fig. 6.**
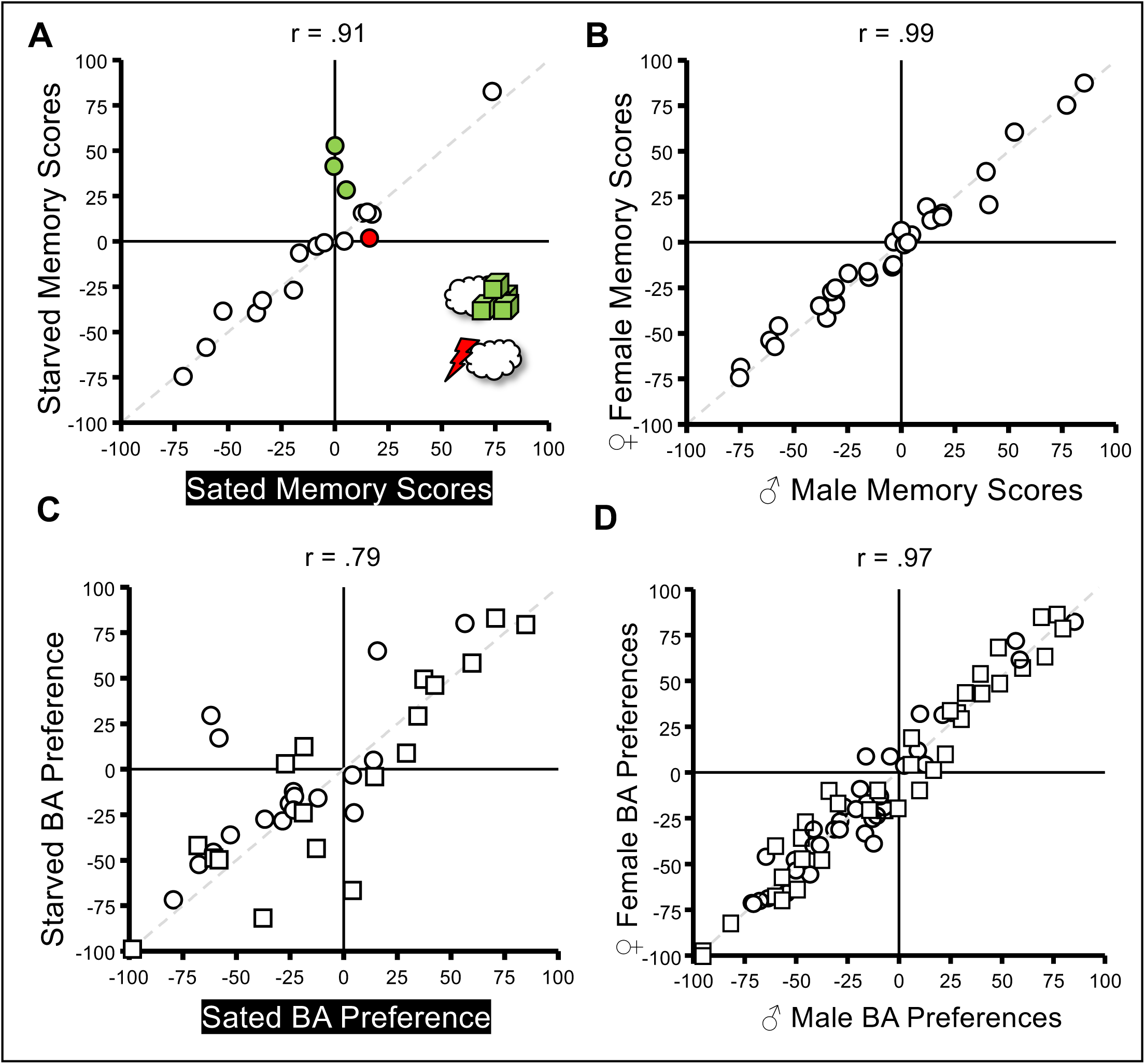
Summary of starvation effects on associative short-term memory in adult flies. **(A)** Correlation plot of memory scores of sated and starved flies across the 18 associative short-term memory tasks used in the present study. Each circle represents the median memory score of the sated/ starved flies for the experiments depicted in Fig. 1, Fig. 2, Fig. 3, Fig. 4, and Fig. 5 (in the case of the odour-D sucrose association, the data from Fig. 1B and Fig. 1C were pooled, as were the data for odour-shock trace conditioning with ISI=-45s from Fig. 3B and Fig. 3C). The dashed grey line represents (y = x), i.e. identity of memory scores in sated and starved flies. Coloured fill emphasizes the exceptional cases, i.e. the increase in memory scores in starved flies in the odour-sugar association task (green) and the decrease in memory scores in starved flies in the shock-odour ’relief’ association task (red). **(B)** As in (A), for the data separated by sex, suggesting no differences in associative short-term memory between female and male flies. **(C)** As in (A), for the odour preferences underlying the memory scores, suggesting no differences in learned odour choice between sated and starved flies. Circles refer to the cases when benzaldehyde (BA) was paired with the reinforcer whereas octanol (OCT) was presented alone; squares refer to reciprocal cases (for plots showing these preferences see Fig. S2, Fig. S3, Fig. S4, Fig. S5, and Fig. S6; in the case of the odour-D sucrose association, the data from Fig. S2A and Fig. S2B were pooled, as were the data for odour-shock trace conditioning with ISI=-45s from Fig. S4A and Fig. S4B). **(D)** As in (B), for the odour preferences underlying the memory scores, suggesting no differences in odour choice between female and male flies (for plots showing these preferences see Fig. S2, Fig. S3, Fig. S4, Fig. S5, and Fig. S6; in the case of the odour-D sucrose association, the data from Fig. S2A’ and Fig. S2B’ were pooled, as were the data for odour-shock trace conditioning with ISI=-45s from Fig. S4A’ and Fig. S4B’). At the top of the panels, ’r’ refers to the Pearson correlation coefficient. The plotted data are reported in the supplemental data file “Starvation DATA”.

### Experiments with larvae

*Drosophila* are insects that undergo full metamorphosis. Adult flies lay their eggs onto ripe to overripe fruit, where, after hatching from the egg, the larvae are mainly concerned with feeding and growth for approximately one week. Then they can leave the food, pupariate and, after approximately one additional week, hatch from the pupal case as adults and renew the cycle of life. During larval stages, tastant cues are thus of outstanding significance to the animals. This has made it possible to develop ecologically valid, robust association tasks for odours and tastants (Gerber and Stocker, 2007; Gerber et al., 2009; Diegelmann et al., 2013; Thum and Gerber, 2019). Here we enquire into the effects of starvation on olfactory associative learning about fructose as a sugar reward (Saumweber et al., 2011; Scherer et al., 2003; Schleyer et al., 2011; Schleyer et al., 2018) and about quinine as a ’bitter’ taste punishment (El-Keredy et al., 2012; Gerber and Hendel, 2006).

### Starvation compromises larval sugar memories

Sated or starved larvae underwent differential conditioning such that one odour was paired with a sugar reward, whereas the other odour was presented unpaired from sugar; this was followed by testing separately for paired- and unpaired memories, taking advantage of a paradigm that assesses these memories relative to baseline odour preference (**Fig. 7A**, **Fig. 7B**). Such a measure of baseline preference is possible because the learned search for sugar, directed towards or away from the odour after paired or unpaired training respectively, adaptively ceases when the sought-for sugar is present during testing (for more details see *Association of odour and sugar reward* in the *Materials and Methods* section). This revealed that both paired and unpaired memories were compromised in starved animals (**Fig. 7C**), whereas baseline odour preferences, cleared of associative effects but reflecting naïve odour preference plus the non-associative effects of handling, odour-exposure, and sugar-exposure, remained unaffected by starvation (**Fig. 7D**).

**Fig. 7.**
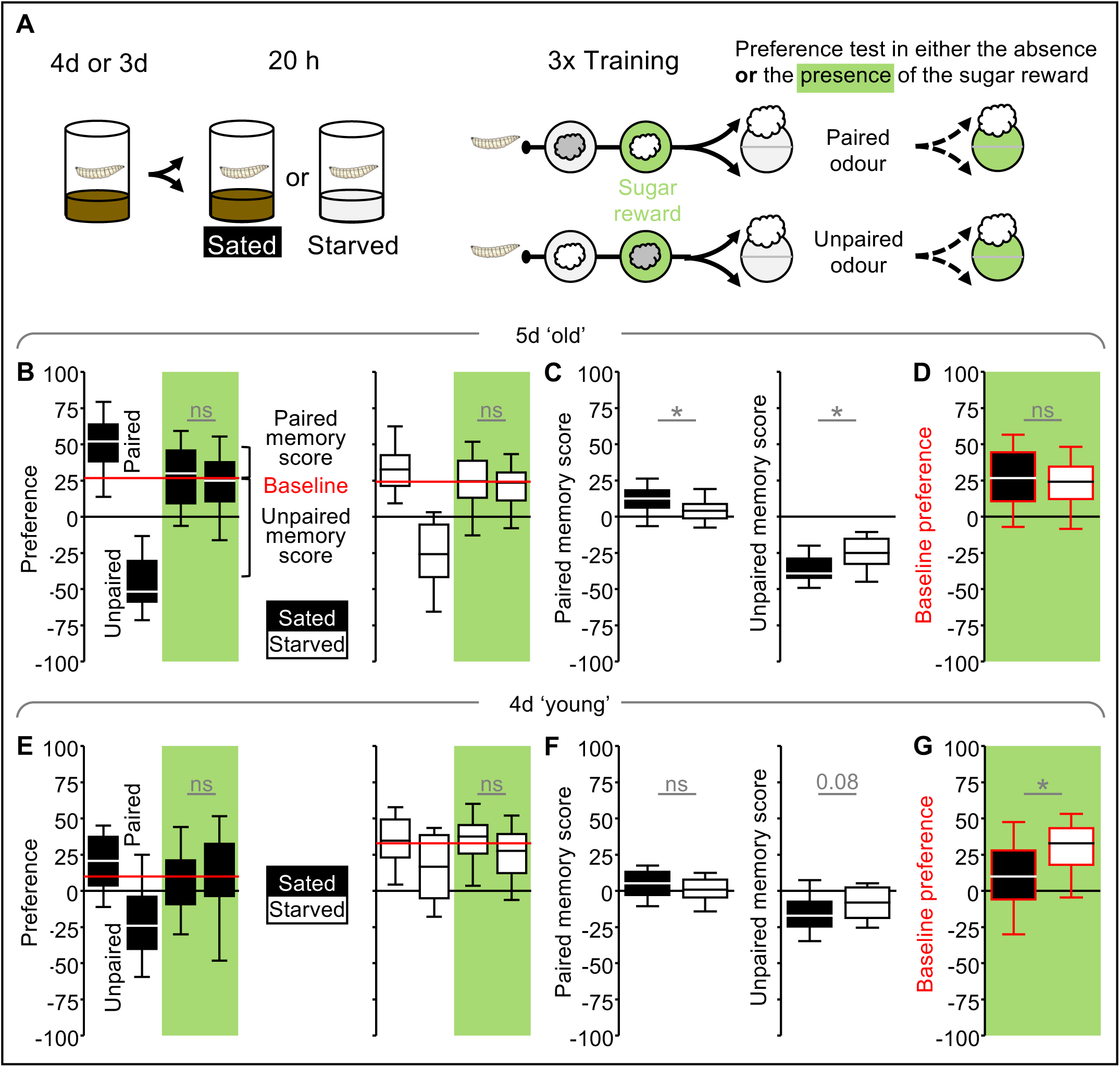
Starvation compromises larval sugar memories. **(A)** Larvae aged 3 or 4 days (d) after egg laying were either kept on their regular food (Sated) or starved for 20 hours (h) (Starved). During training, one of two odours (grey and white clouds) was presented on a plain agarose Petri dish (white circle), whereas the other odour was presented with fructose as a reward in addition (green circle). Three such training trials were followed by a preference test for the odour that during training had been presented paired with or unpaired from the reward. The larvae on the odour side and the blank side of the Petri dish were counted, and odour preferences were calculated according to Eqn 4, with positive and negative scores indicating a preference for and avoidance of the odour, respectively. Testing was carried out either on a plain agarose Petri dish or on a Petri dish additionally containing the fructose reward; the latter allows baseline odour preferences cleared of associative effects to be measured. The chemical identity of the training odours was balanced across repetitions of the experiment; the identity of the testing odour was kept the same throughout. For the larvae already collected 3 days after egg laying, smaller, 55mm-diameter Petri dishes were used to accommodate the smaller size of these animals. **(B)** Odour preferences of sated (filled boxplots) and starved (open boxplots) larvae. Odour preferences were higher than baseline if the odour had been paired with reward, and below baseline if the odour had been presented unpaired from the reward. The baseline odour preference (red line) was calculated, separately for sated and starved larvae, as the median of the pooled odour preferences of the larvae tested in the presence of the reward (MWU tests, for sated and starved larvae: U=687.5, p=0.47, N=39, 39; U=653.5, p=0.48, N=38, 38). **(C)** Based on the odour preferences in (B), paired memory scores and unpaired memory scores were calculated according to Eqn 5 and Eqn 6. Relative to sated larvae (filled boxplot), both paired and unpaired memory scores were compromised in starved larvae (open boxplot) (MWU tests, for paired and unpaired memory scores: U=462, p<0.05, N=39, 38; U=395, p<0.05, N=39, 38). **(D)** Baseline odour preferences do not differ between sated and starved larvae (MWU test: U=2657.5, p=0.27, N=78, 76). Boxplots represent data pooled from the respective odour preferences in (B). **(E-G)** As in (B-D), for larvae collected aged 3 days (d) after egg laying (E: MWU tests, for sated and starved larvae: U=695, p=0.5159, N=39, 39; U=577.5, p=0.0682, N=39, 39) (F: MWU tests, for paired and unpaired memory scores: U=596, p=0.10, N=39, 39; U=587, p=0.08, N=39, 39) (G: MWU test: U=1747, p<0.05, N=78, 78). *indicates significance in MWU tests at p<0.05, and “ns” non-significance in these tests, at p>0.05. Statistical results, including OSS tests of memory scores against chance levels (i.e. zero), are reported along with the source data in the supplemental data file “Starvation DATA”.

Given the improvements in sugar memory upon starvation in adult flies (**Fig. 1**), we were surprised by these impairments in the case of larvae and wondered whether starvation might accelerate the larval-pupal transition. That is, once larvae have passed the critical weight checkpoint during the third larval instar, they might take a developmental ’emergency exit’ from their larval lives and transition into an early pupa (Shingleton et al., 2005). As part of the regular larva-to-pupa transition, the mushroom bodies undergo a complex metamorphic rearrangement, effectively disrupting DAN-KC-MBON connectivity (Truman et al., 2023). We reasoned that if starvation were to prompt a developmental trajectory into early pupariation, a disruption in mushroom body connectivity might be prompted too, and this might in turn compromise mnemonic function. We therefore hypothesized that sugar memories should not be compromised when the starvation period starts before an early pupariation is an option for the larvae, i.e. when they are 3 days old (**Fig. 7A**). Although technically speaking this is the case (**Fig. 7E**, **Fig. 7F**), we note that in these younger animals neither sated nor starved larvae showed evidence of paired memory to begin with (**Fig. 7F**, **left**), and that there is an apparent trend for diminished unpaired memory in these younger animals that indeed runs against our hypothesis (p= 0.08) (**Fig. F, right**). We further note that in these younger larvae starvation leads to an increase in baseline odour preferences (**Fig. 7G**), possibly reflecting starvation-induced differences in naïve odour preference (but see *Innate preference behaviour* section below) and/or differences with respect to non-associative effects of handling, of odour-exposure or of sugar-exposure that are averaged out when associative memory scores are calculated (see *Discussion*).

### Starvation compromises larval quinine memories

We next repeated the above experiments using quinine as a bitter taste punishment (**Fig. 8**). In this case too, a measure of baseline preferences is possible because an escape from quinine, directed away from or towards the odour after paired or unpaired training respectively, is adaptive only when escape is warranted by the presence of quinine during testing (for more details see *Association of odour and bitter-taste punishment* in the *Materials and Methods* section). In comparison to the sated condition, starvation of 4-day-old larvae tendentially diminished paired quinine memory (p= 0.047) (**Fig. 8B**, **Fig. 8C, left**) and compromised unpaired memory (**Fig. 8B**, **Fig. 8C, right**). We furthermore observed a higher baseline odour preference in starved larvae than in sated animals (**Fig. 8D**).

**Fig. 8.**
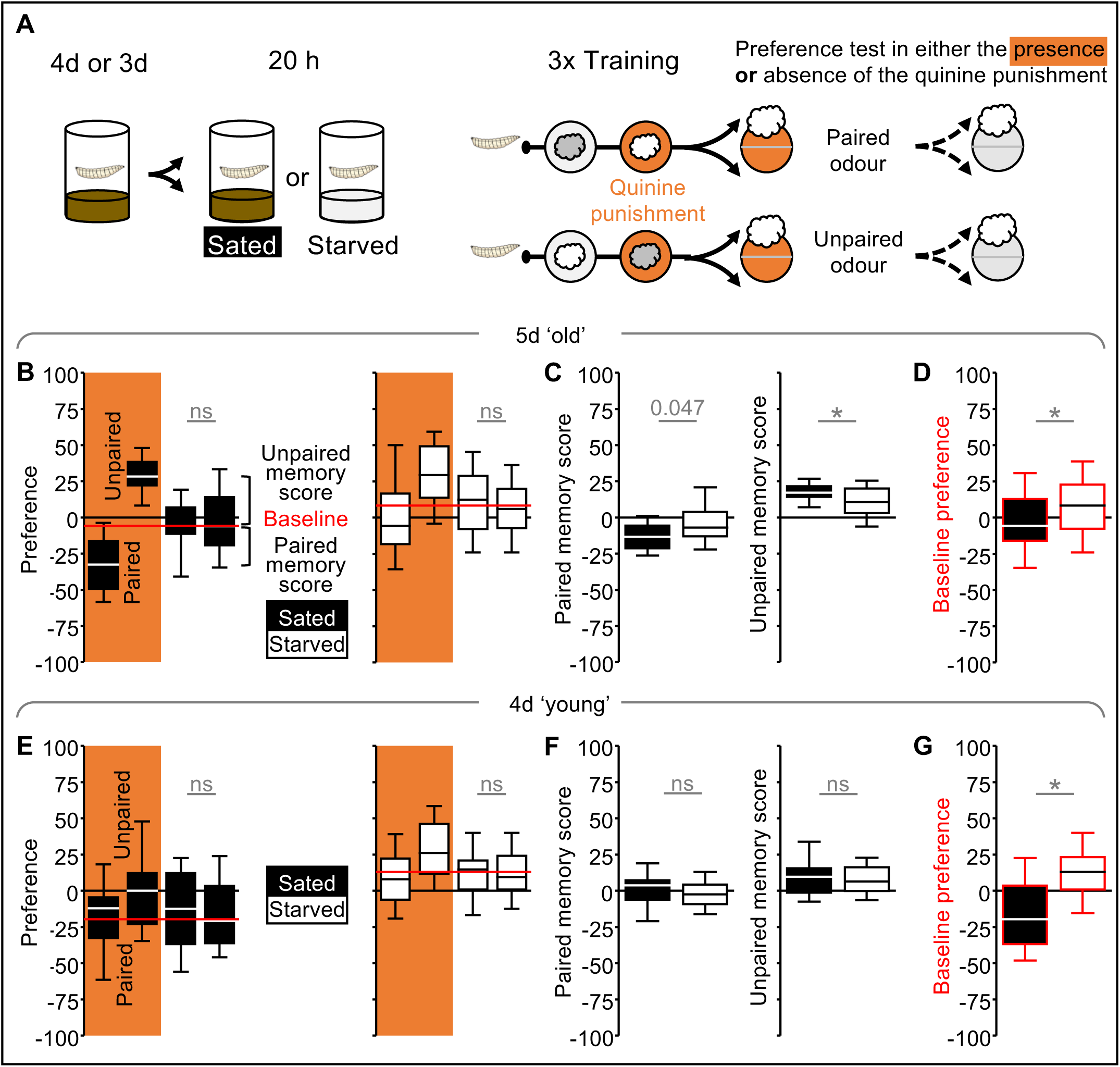
Starvation compromises larval quinine memories. **(A)** Larvae aged 3 or 4 days (d) after egg laying were either kept on their regular food (Sated) or starved for 20 hours (h) (Starved). During training, one of two odours (grey and white clouds) was presented on a plain agarose Petri dish (white circle) whereas the other odour was presented with quinine as a taste punishment in addition (orange circle). Three such training trials were followed by a preference test for the odour that during training had been presented paired with or unpaired from the taste punishment. The larvae on the odour side and the blank side of the Petri dish were counted, and odour preferences were calculated according to Eqn 4, with positive and negative scores indicating a preference for and avoidance of the odour, respectively. Testing was carried out either on an agarose Petri dish additionally containing the quinine punishment or on a plain agarose Petri dish; the latter allows baseline odour preferences cleared of associative effects to be measured. The chemical identity of the training odours was balanced across repetitions of the experiment; the identity of the testing odour was kept the same throughout. For the larvae already collected 3 days after egg laying, smaller, 55mm-diameter Petri dishes were used to accommodate the smaller size of these animals. **(B)** Odour preferences of sated (filled boxplots) and starved (open boxplots) larvae. Odour preferences were below baseline if the odour had been paired with punishment, and higher than baseline if the odour had been presented unpaired from the punishment. The baseline odour preference (red line) was calculated, separately for sated and starved larvae, as the median of the pooled odour preferences of the larvae tested on a plain agarose Petri dish (MWU tests, for sated and starved larvae: U=331, p=0.91, N=26, 26; U=302.5, p=0.52, N=26, 26). **(C)** Based on the odour preferences in (B), paired memory scores and unpaired memory scores were calculated according to Eqn 5 and Eqn 6. Relative to sated larvae (filled boxplots), paired memory scores were tendentially reduced (MWU tests: U=229, p=0.047, N=26, 26) and unpaired memory scores compromised (MWU tests: U=214, p=0.0238, N=26, 26) in starved larvae (open boxplots). **(D)** Relative to sated larvae, baseline odour preferences were increased in starved larvae (MWU test: U=966, p=0.01, N=52, 52). Boxplots represent data pooled from the respective odour preferences in (B). **(E-G)** As in (B-D), for larvae collected aged 3 days (d) after egg laying (E: KW test p=0.08, N=37, 37; U=678, p=0.95, N=37, 37), (F: MWU tests, for paired and unpaired memory scores: U=547, p=0.14, N=37, 37; U=632, p=0.57, N=37, 37), (G: MWU test: U=1230.5, p<0.05, N=74, 74). * indicates significance in an MWU test, at p<0.05, and “ns” indicates non-significance in an MWU test, at p>0.05. Statistical results, including OSS tests of memory scores against chance levels (i.e. zero), are reported along with the source data in the supplemental data file “Starvation DATA”.

In 3-day-old larvae, both paired and unpaired memory scores were relatively small, and no decreases induced by starvation were detectable (**Fig. 8E**, **Fig. 8F**). This contrasts with a pronounced increase in baseline odour preferences in these younger larvae (**Fig. 8G**).

As a summary of the results in larvae, we plotted the median associative memory scores of starved versus sated animals (**Fig. 9A**), suggesting that relative to the sated condition, memory scores tend to be generally diminished upon starvation. Analysis of the odour preferences underlying these memory scores furthermore suggests that these odour preferences are generally increased upon starvation (**Fig. 9B**), possibly reflecting non-associative effects of handling, of odour-exposure or of sugar-exposure (see *Discussion*).

**Fig. 9.**
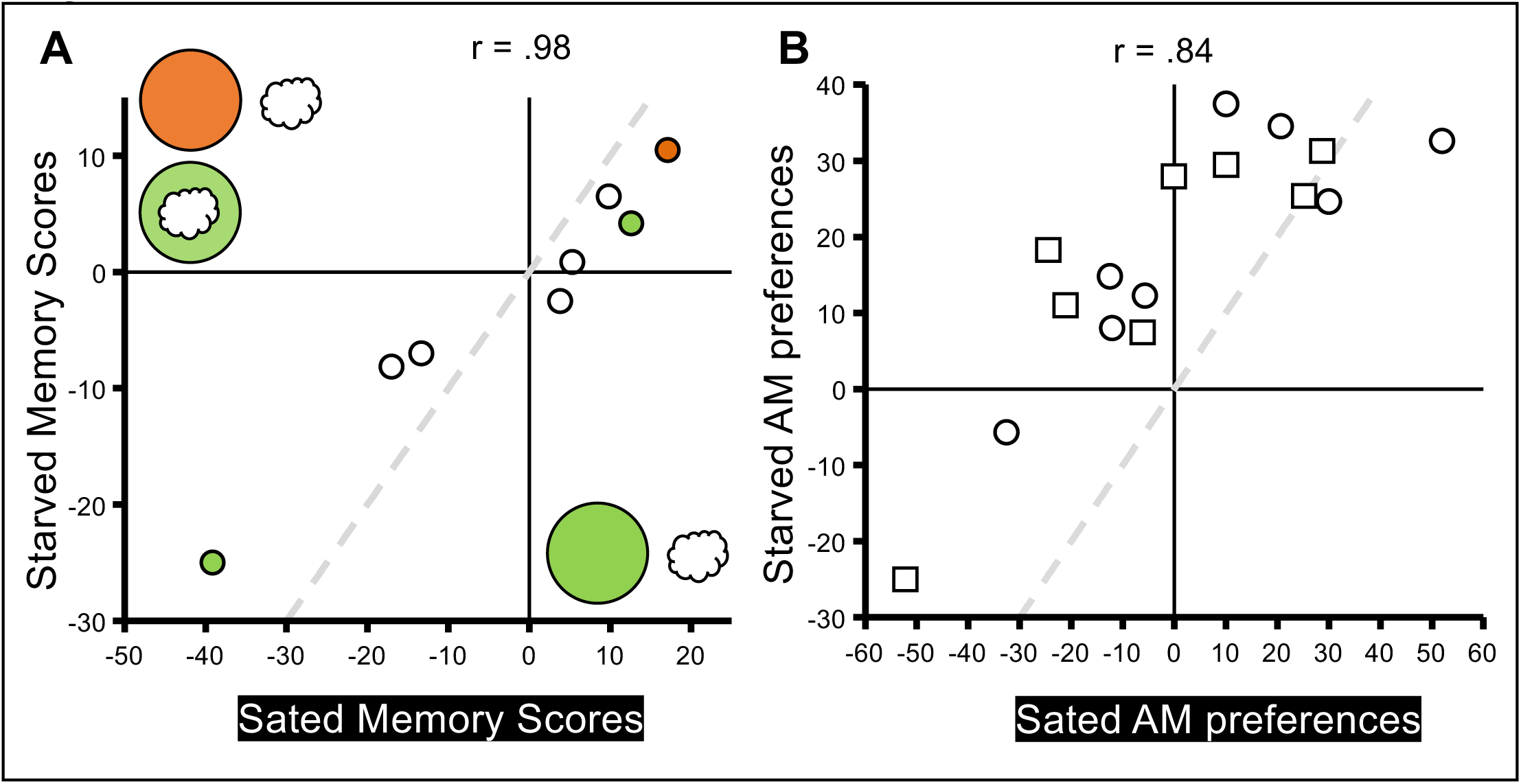
Summary of starvation effects on associative short-term memory in larvae. **(A)** Correlation plot of memory scores of sated and starved larvae across the eight association tasks used in the present study. Each circle represents the median of the paired or unpaired memory score of the sated/ starved larvae for the experiments depicted in Fig. 7 and Fig. 8. The dashed grey line represents (y = x), i.e. identity of memory scores in sated and starved larvae. Though coloured fills emphasize the significantly affected memory scores in the indicated tasks, we note that relative to the sated condition memory scores appear generally diminished upon starvation. **(B)** As in (A), for the odour preferences underlying the memory scores, suggesting a general increase in odour preference upon starvation. Circles refer to the cases when n-amyl acetate (AM) was paired with the reinforcer, whereas squares refer to presentations of AM unpaired from the reinforcer. At the top of the panels, ’r’ refers to the Pearson correlation coefficient. The plotted data are reported in the supplemental data file “Starvation DATA”.

### Innate preference behaviour and larval locomotion

We wondered not only whether starvation might affect associative memory but also how the stimuli involved are processed ’innately’, i.e. in experimentally naïve animals, and whether starvation might lead to impairments in task-relevant motor abilities. As no effect of starvation was observed in adult flies in any of the odour-shock association tasks (**Fig. 2B, left**, **Fig. 3**, **Fig. 5**), the task-relevant processing of odours and shock, as well as movement faculties, are apparently intact. However, as previously reported (Coban et al., 2024; Gruber et al., 2013), starvation leads to an increase in ’innate’ sugar preference in adult flies (**Fig. 10A**).

**Fig. 10.**
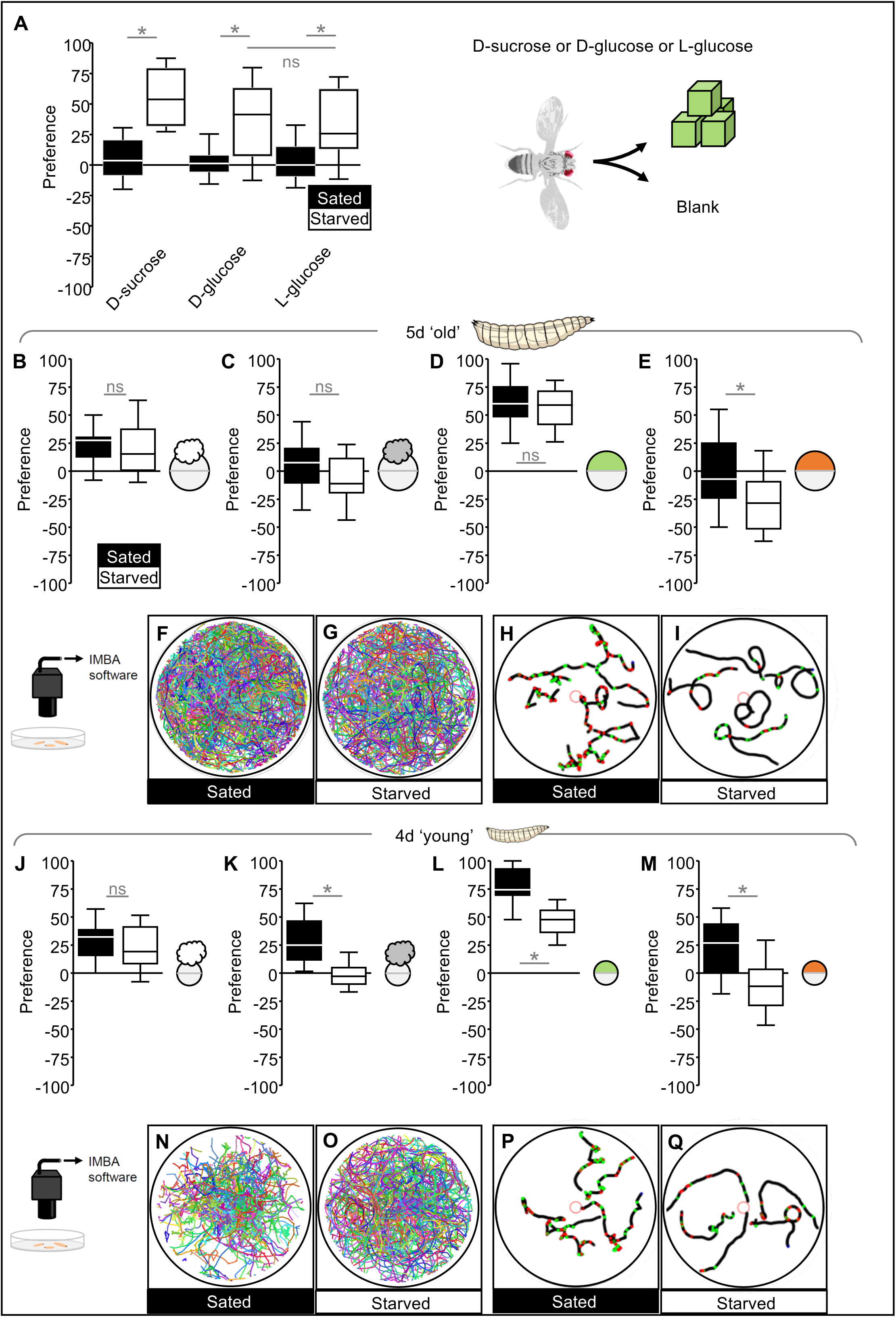
Effects of starvation on innate preferences and larval locomotion. **(A)** Using a starvation protocol as in Fig. 1, cohorts of ∼30 sated or starved experimentally naïve wild-type flies were tested for their preference for D-sucrose, D-glucose or L-glucose (green cubes) in a T-maze. Based on the number of flies in each arm of the maze, preferences were calculated with due modification of Eqn. 4, but with the same logic, with positive and negative scores indicating preference for and avoidance of sugar, respectively. Relative to the sated condition, preferences for all three sugars were increased upon starvation (MWU-tests sated vs. starved for D-sucrose: U=290, p<0.05, N= 60, 60; for D-glucose: U=751, p<0.05, N=60, 60; for L-glucose: U=783.5, p<0.05, N=60, 60. MWU-test starved D-glucose vs. starved L-Glucose: U=1739, p=0.75, N=60, 60). **(B-E)** Using a starvation protocol as in Fig. 7, cohorts of ∼20 ’old’ sated or starved larvae were tested for their preference for n-amyl acetate (white cloud) (B), octanol (grey cloud) (C), fructose (green) (D), or quinine (orange) (E) in a Petri dish assay (circle). Based on the number of larvae on each side of the Petri dish, preferences were calculated with due modification of Eqn. 4, but with the same logic, with positive and negative scores indicating preference for and avoidance of the respective stimulus. Relative to the sated condition, preferences in starved larvae were unchanged for both odours and for fructose, whereas quinine became more aversive upon starvation (MWU-tests, for n-amyl acetate: U=144, p=0.58, N=18, 18; for octanol: U=112.5, p=0.28, N=17, 17; for fructose: U=256.5, p=0.70, N=22-25; for quinine: U=1195.5, p<0.05, N=64, 62). **(F-I)** Using a starvation protocol as in Fig. 7, cohorts of 7-8 5-day ’old’ sated or starved larvae were placed on a Petri dish, and their behaviour was videorecorded for 3 minutes. To illustrate the animals’ behaviour, we plotted all the tracks of all recorded (F) sated and (G) starved larvae (663 and 439 tracks, respectively), as well as five randomly selected sample tracks of (H) sated and (I) starved larvae. The starved larvae apparently make fewer lateral head movements, so-called head casts (red and green dots). For a more detailed analysis, see Fig. S8. **(J-M)** As in (B-E), for ’young’ larvae, using smaller, 55-mm Petri dishes to accommodate the smaller size of these animals. Relative to the sated condition, preferences for n-amyl acetate were tendentially decreased (J); octanol (K) and fructose (L) preferences were reduced; and quinine became more aversive (M) upon starvation (MWU-tests, for n-amyl acetate: U=173.5, p=0.48, N=20, 20; for octanol: U=46.5, p<0.05, N=20, 20; for fructose: U=65, p<0.05, N=24, 24; for quinine: U=409.5, p<0.05, N=45, 45). **(N-Q)** As in (F-I), for ’young’ larvae (N: 290 tracks; O: 408 tracks), which also make fewer head casts when starved (red and green dots). For a more detailed analysis, see Fig. S8. * indicates significance in MWU tests at p<0.05, and “ns” non-significance in these tests, at p>0.05. Statistical results, including OSS tests against chance levels (i.e. zero), are reported along with the source data in the supplemental data file “Starvation DATA”.

In contrast, in 5-day-old larvae, ’innate’ preferences for odours and sugar are unaffected by previous starvation (**Fig. 10B**, **Fig. 10C**, **Fig. 10D**), whereas ’innate’ behaviour in relation to quinine becomes more aversive (**Fig. 10E**). In addition, we compared larval locomotion in the absence of any directional stimuli between sated and starved animals (**Fig. 10F**, **Fig. 10G**). In comparison to sated larvae, starved larvae appeared to perform fewer lateral head movements, so-called head casts (**Fig. 10H**, **Fig. 10I**), giving a less zigzagging appearance to their tracks. A more detailed analysis, normalized to individual body lengths, confirmed that starved larvae did indeed move faster in a forward direction and less laterally, and that their peristaltic forward movement was characterized by larger yet longer-lasting ’steps’ (**Fig. S8**).

The situation is again slightly different in younger, 4-day-old larvae, as in these animals ’innate’ preferences generally decrease upon starvation (**Fig. 10J**, **Fig. 10K**, **Fig. 10L**, **Fig. 10M**). Regarding their locomotion, the differences between sated and starved larvae generally matched those of their older counterparts, with faster forward movement, less lateral movement, and larger, but in this case slightly less long-lasting, peristaltic steps upon starvation, bestowing a less zigzagging appearance on the tracks of 4-day-old larvae too (**Fig. 10N**, **Fig. 10O**, **Fig. 10P**, **Fig. 10Q, Fig. S8**).

## Discussion

This study examined the effects of starvation on memory in *Drosophila melanogaster* across 26 different tasks. Memory scores remained unaffected in 19 of the cases, were impaired in four of them, and improved in three cases. On the one hand, such task-dependence is to be expected, given that the tasks differ in a number of respects:

- the reinforcers used (different kinds of sugar, electric shock, optogenetic activation of the PPL1-01 DANs, or quinine);
- the number of training trials (1, 3, or 6);
- the life stage and age of the subjects (2-day to 5-day-old adult flies, 4-day or 5-day-old larvae);
- the predictive structure and associative timing of the tasks (odours associated with the presence, absence, the occurrence, or the termination of the reinforcer);
- whether memory is expressed as an increase or as a decrease in odour preference; and
- whether learned behaviour is motivated by pursuing reward or by avoiding/ escaping punishment.

On the other hand, the observed task-dependence is surprising, given that all the tasks address associative short-term memory, involve odours as predictive cues, use differential classical conditioning, and employ behavioural testing in choice situations. This encompasses only a narrow subset of the mnemonic faculties that have been reported in flies, which include sensitization and habituation as non-associative forms of memory, latent learning, non-differential forms of classical conditioning, operant conditioning, medium- and long-term memory, and learning about cues from sensory modalities other than olfaction, or from social interactions. Obviously, the current findings cannot be extrapolated to any of these forms of learning. Also, assessing variables other than choice, or parametric variations in, for example, the duration or the developmental timing of starvation, may well paint a picture different from what we report (adult flies: **Fig. 6**; larvae: **Fig. 9**).

### Does the ’adaptive specificity hypothesis’ apply in adult flies?

Our results for adult flies support the ’adaptive specificity hypothesis’, the notion that starvation specifically enhances food-related processing (Dethier, 1976; Kim et al., 2017; Sakagiannis et al., 2025; Selcho and Pauls, 2019; Sgammeglia and Sprecher, 2022; Suarez-Grimalt et al., 2024). This is because the only case of improved memory is for odour-sugar associations (**Fig. 1**) (**Fig. 6**). In contrast, all tasks yielding negative memory scores are unaffected and, strikingly, tasks that – just like odour-sugar associations – also yield positive memory scores, namely relief associations and extinction memory, are either unaffected too or even impaired (**Fig. 2**, **Fig. 4**, **Fig. 5**) (**Fig. 6**). Thus, it is when the task is motivated by the pursuit of a sugar reward that memory is improved, whereas this is not the case for tasks that are otherwise motivated. In terms of the mechanisms underlying this improvement, we note that downstream of the primary internal ’hunger sensors’ and ’satiation signals’ a complex reconfiguration of the biogenic amine systems effectuates state-dependent adaptations (Kim et al., 2017; Selcho and Pauls, 2019; Suarez-Grimalt et al., 2024). Starvation leads to a decrease in the activity of octopaminergic/tyraminergic OA-VL neurons, reducing the sensitivity of bitter sensory neurons (LeDue et al., 2016). In turn, promoting activity in the likewise octopaminergic/tyraminergic OA-VPM4 neurons (Youn et al., 2018) and starvation-induced dopamine signalling from TH-VUM neurons (Marella et al., 2012) can increase the sensitivity of sugar sensory neurons, suggesting an aminergic push-pull mechanism by which starvation inflates the valuation of gustatory input (see also Inagaki et al., 2014). Fittingly, in starved flies, signalling from octopaminergic/tyraminergic neurons (TDC-Gal4) is required for low-quality sugars to be rewarding (Burke et al., 2012) (for a similar phenotype in TβH mutants: Berger et al., 2024, *loc. cit.* Figure 1). These effects are brought about at least partially by signalling to rewarding DANs in the horizontal lobes (Burke et al., 2012; Huetteroth et al., 2015).

Our starvation protocol improves odour-sugar memories mediated by both the ’sweetness’ and the metabolic value of the sugar (**Fig. 1C**): the observation that in the starved state, memory scores for D-glucose are higher than for L-glucose suggests that starvation promotes a memory component that is based on the higher metabolic value of D-glucose. In turn, the fact that in the starved state the flies form a memory for L-glucose at all suggests that starvation also promotes a memory component based on the sensory characteristics of glucose (its ’sweetness’). Our interpretation of why starvation improves even a metabolically worthless ’sweetness’ memory is that this supports a working-memory-like process to ’bridge the gap’ to learning about sugar-predictors even at a moment when the metabolic feedback from sugar ingestion cannot yet have arrived.

Whereas our own data and the available literature thus consistently show that starvation improves odour-sugar associations, there is conflicting evidence with respect to odour-shock associations. Our data (**Fig. 1**), as well as data from Ichinose and Tanimoto (2016) and from Coban et al. (2024), show the above-mentioned improvement in odour-sugar association, together with either no effect (**Fig. 2**, **Fig. 3**, **Fig. 5**) (Ichinose and Tanimoto, 2016) or even an impairment (Coban et al., 2024) in odour-shock associations, confirming the ’adaptive specificity hypothesis’. However, Gruber et al. (2013) showed that both odour-sugar and odour-shock associations are improved by starvation, with the latter result confirmed by Meschi et al. (2024), challenging the ’adaptive specificity hypothesis’. To the best of our judgement, variations in procedure between these studies are minor, suggesting that the boundary conditions under which starvation improves, impairs, or has no effect on odour-shock associations still need to be identified (Strack, 2017). We note that starvation-induced octopamine/tyramine signalling towards rewarding DANs in the lobes may specifically promote odour-sugar associations (see preceding paragraph), whereas octopamine/tyramine signalling in the mushroom bodies’ input region, the calyx, may provide a ’salience signal’ that promotes association formation more generally (Rudelt et al., 2025).

### Why does starvation not affect PPL1-01 reinforcement?

Starvation does not affect associations of odour and optogenetic activation of PPL1-01 DANs, whether in trace, delay or relief conditioning (**Fig. 4**). This is consistent with Liu et al. (2012), who showed that delay conditioning with thermogenetic activation of a broader set of DANs including PPL1-01 (TH-Gal4) is not affected by starvation either. Both these observations are surprising, however, given reports of starvation-induced changes in PPL1-01 physiology. Using the NP0047-Gal4 driver strain covering PPL1-01 plus PPL1-03, Placais and Preat (2013) observed that starvation dampens spontaneous calcium oscillations in these cells (*loc. cit.* Figure S3). Furthermore, Meschi et al. (2024) used the same driver strain as in the present study (MB320C-Gal4, in their case combined with the transcriptional calcium reporter CaLexA), and showed that starvation decreases spontaneous activity in PPL1-01; using a different driver strain (MB504B-Gal4, combined with the dopamine sensor GRAB_DA2m_), the authors also showed that starvation reduces spontaneous dopamine release from PPL1-01, but enhances shock-evoked dopamine release. Concerning the present study, one may argue that this syndrome of starvation-induced changes may be overridden by strong opto- (**Fig. 4**) or thermogenetic (Liu et al., 2012) activation of PPL1-01. Such an override of starvation-induced changes in PPL1-01 physiology may also be the reason why PPL1-01 relief remains intact upon starvation (**Fig. 4**), whereas shock-relief is impaired (**Fig. 2**). Alternatively, starvation may affect neurons other than PPL1-01 to impair shock-relief.

### Can the ’adaptive specificity hypothesis’ be confirmed in larvae?

Our data for larvae do not support the ’adaptive specificity hypothesis’: memories for both paired and unpaired presentations of odour and sugar, both of which serve the search for sugar and thus are appetitively motivated, are compromised by starvation, rather than improved by it, in 5-day-old animals (**Fig. 7**). To varying degrees, similar detrimental effects are observed in larvae starved earlier in life and also for aversively motivated quinine associations (**Fig. 8**, **Fig. 9**). When testing for memory immediately after single-trial conditioning, Brünner et al. (2020) likewise found that odour-sugar associations were impaired upon starvation, and when three training trials were used, as in the present study, a corresponding tendency was observed. Thus, to the extent that starvation affects memory in larvae it does so with opposite sign and more generally than the ’adaptive specificity hypothesis’ suggests.

Considering longer-lasting memories, Brünner et al. (2020) observed an improvement in memory when the larvae were tested 80 min after three-trial conditioning. Such an improvement was still observed in a mutant of the *dunce* gene known to be impaired in what is called the amnesia-sensitive memory component (ASM) but intact in the amnesia-resistant memory component (ARM). In turn, starvation did not increase memory in a mutant of the *radish* gene that lacks that ARM component, suggesting that it is ARM that is promoted by starvation, a conclusion similar to what was established for longer-lasting aversive memories in adult flies (Basu et al., 2024; Placais and Preat, 2013).

### Implications for DAN function in larvae?

The dopaminergic neuron DAN-h1 has not yet differentiated in first-instar larvae (Saumweber et al., 2018), and it is unknown whether DAN-h1 is already functional in the younger animals tested here (**Fig. 7E**, **Fig. 7F**, **Fig. 7G**, **Fig. 8E**, **Fig. 8F**, **Fig. 8G**). If it were not, this could account for the absence of odour-sugar paired memory in the sated group (**Fig. 7F, left**), given that blocking DAN-h1 activity reduces odour-sugar memory scores to half in a paradigm that measures the combined effects of paired and unpaired training (Saumweber et al., 2018).

We further note that DAN-g1 has been implicated in non-associative learning (Saumweber et al., 2018, *loc.cit.* Suppl. Fig. 8). Specifically, Saumweber et al. (2018) reported that pre-exposure to sugar non-associatively increases subsequent odour preferences – and that this increase is further enhanced when DAN-g1 activity is blocked. It would be interesting to test whether starvation compromises DAN-g1 activity, contributing to the non-associative increases in odour preference observed in the present study (**Fig. 9**) or, as recent results from Weber et al. (2025) may suggest, to compromised aversive memory upon starvation.

### Effects of starvation on innate preferences

Starvation may affect not only associative memory but also the way the stimuli to be associated are processed ’innately’, i.e. in experimentally naïve animals, or the faculties of movement required in a task. For adult flies the task-relevant processing of odours and shock, as well as movement faculties, apparently remain intact, because in none of the odour-shock association tasks was an effect of starvation observed (**Fig. 2B, left, Fig. 3**, **Fig. 5**), a conclusion consistent with a lack of effect of starvation on odour- or shock processing noted in previous reports (Coban et al., 2024; Meschi et al., 2024). We did, however, observe that starvation leads to an increase in ’innate’ sugar preference (**Fig. 10A**), in line with previous studies (Coban et al., 2024; Gruber et al., 2013). Although the neuronal pathways underlying innate sugar preference and the rewarding effects of sugar certainly do not overlap completely, it is plausible that the starvation-induced improvement in odour-sugar association (**Fig. 1**) (Coban et al., 2024; Gruber et al., 2013; Ichinose and Tanimoto, 2016; Tempel et al., 1983) is based at least partially on enhanced sensory processing of sugar.

In 5-day-old larvae, preferences for odours and sugar are unaffected by starvation (**Fig. 10B**, **Fig. 10C**, **Fig. 10D**), and no change in baseline odour preference is observed (**Fig. 7D**). This suggests that task-relevant alterations in sensory-motor faculties or changes in the non-associative effects of handling or stimulus-exposure cannot account for the compromised odour-sugar associations upon starvation (**Fig. 7B**, **Fig 7C**). This is in line with what Brünner et al. (2020) found regarding the preferences for sugar and for n-amyl acetate, the testing odour used in the present study (for benzaldehyde, an odour not used in the present context, they found a starvation-induced decrease in preference).

The situation is different for younger, 4-day-old larvae, and, regardless of larval age, when quinine is involved. Specifically, we observed strong starvation-induced increases in baseline odour preference (**Fig. 7G**, **Fig. 8D**, **Fig. 8G**), decreases in ’innate’ odour- and sugar-preference (tendentially in **Fig. 10J**), (**Fig. 10K**, **Fig. 10L**) and stronger aversion to quinine (**Fig. 10E**, **Fig. 10M**). This syndrome of changes cannot be explained through starvation-induced motor impairments and is not compatible with the ’adaptive specificity hypothesis’. Following leads from the pioneering work of de Tredern et al. (2025) and Vogt et al. (2021), future work may focus on how starvation affects ’innate’ preferences depending on when and for how long the larvae are starved, and whether they are assayed in groups, as in the present study, or one at a time (Mudunuri et al., 2026). Regarding the strong starvation-induced increases in baseline odour preference that we observed (**Fig. 7G**, **Fig. 8D**, **Fig. 8G**), it seems warranted to investigate whether these come about by changes in the non-associative effects of handling, of odour- or tastant exposure, or of any combination of these.

### Effects of starvation on locomotion

Finally, starvation can change patterns of locomotion in the absence of directional stimuli. In larvae of either age, we found that relative to the sated condition starvation leads to a less zigzagging appearance in locomotor tracks, specifically to increased forward movement and decreased lateral head movement (**Fig. 10H**, **Fig. 10I**, **Fig. 10P**, **Fig. 10Q, Fig. S8**). These behavioural patterns may reflect different strategies of food search: sated larvae may perform a local search as they have only just been removed from food and may expect to be still close to it, whereas starved larvae may rather perform a global search after being starved for almost a day, not expecting to be anywhere close to a food source. Though previous studies have found that such foraging strategies can be genetically determined by a natural polymorphism in the *foraging* gene (Sokolowski, 1980; Anreiter and Sokolowski, 2019), it remains to be tested whether *foraging* function is involved in any of the starvation effects that we have observed in the current study.

In summary, our results are not readily compatible with an interpretation solely according to the ’adaptive specificity hypothesis’ and call rather for case-by-case analyses of whether and how a given learning task, or behaviour, is affected by starvation in *Drosophila*.

## Acknowledgements

Discussions with Katrin Vogt and Claire Eschbach are gratefully acknowledged. We thank Rupert D.V. Glasgow, Zaragoza, Spain, for corrections of earlier drafts of the manuscript, and Bettina Kracht and Melissa Prothmann, Diana Walther, Canan Nazarov, Isabel Walther for experimental contributions and technical assistance.

## Author contributions

Conceptualization: E.S., S.K., B.G.; Data curation: E.S., T.N., M.S., C.K., J.T.; Formal analysis: E.S., S.K., M.S., C.K.; Funding acquisition: A.I.G., M.S., B.G.; Investigation: E.S., S.K., M.B., A.C., S.D., A.I.G., T.N., C.K., J.T.; Project administration: E.S., B.G.; Software: M.T., M.S.; Supervision: B.G.; Validation: all authors. Visualization: E.S., M.T., M.S., C.K., J.T., B.G.; Writing - original draft: E.S., S.K., M.S.; Writing - review & editing: all authors.

---

**Fig. S1.**
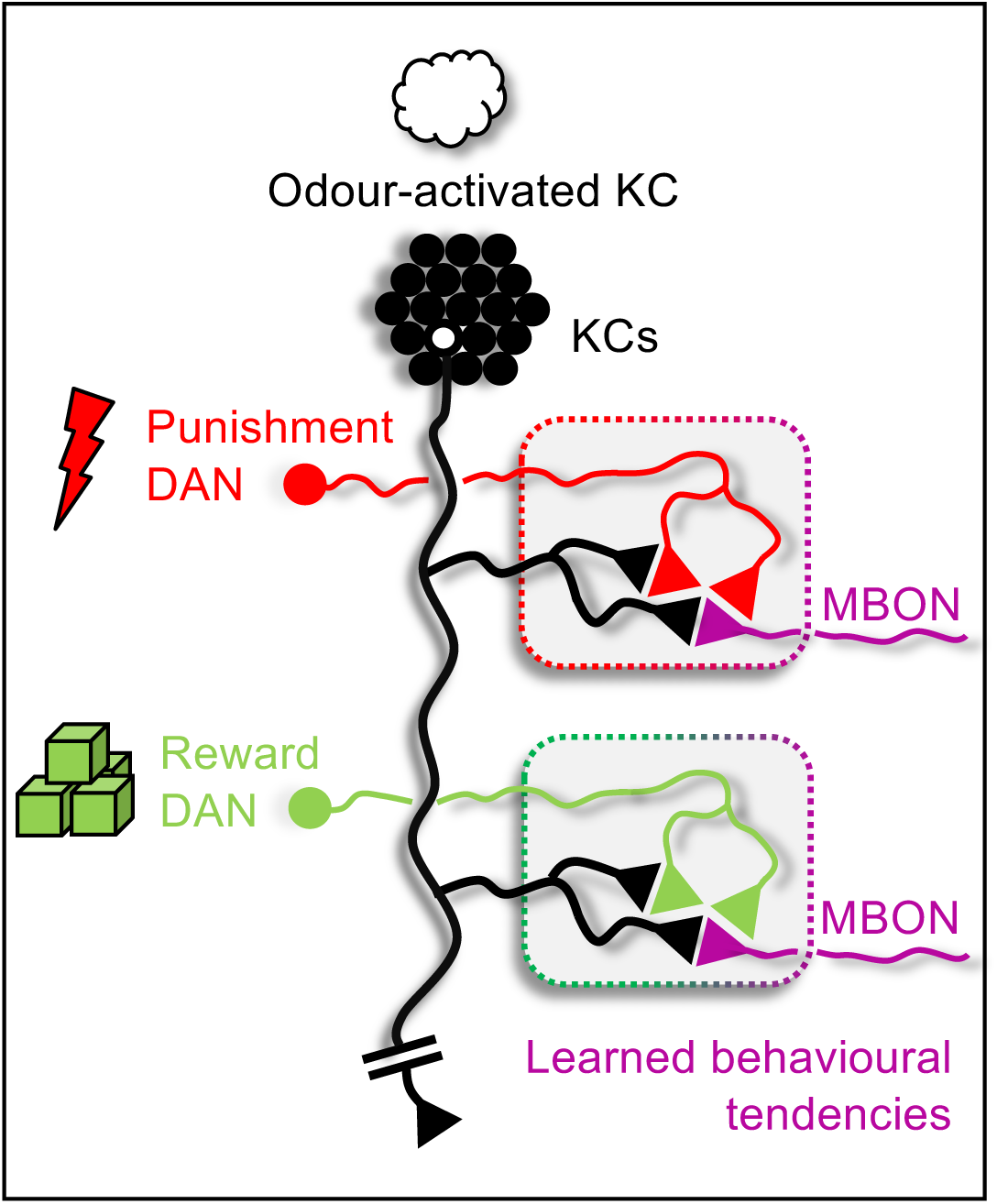
Highly simplified circuit overview of associative learning in the *Drosophila* mushroom body. An odour (white cloud) activates a sparse population of Kenyon cells (KCs), which project to mushroom body output neurons (MBONs, magenta colour). Dopaminergic neurons (DANs) convey punishment (red) or reward (green) signals that modulate KC::MBON synapses within distinct compartments. Coincidence of odour-evoked KC activity and DAN input induces synaptic plasticity, leading to changes in MBON output that bias behavioural tendencies toward avoidance or approach.

**Fig. S2.**
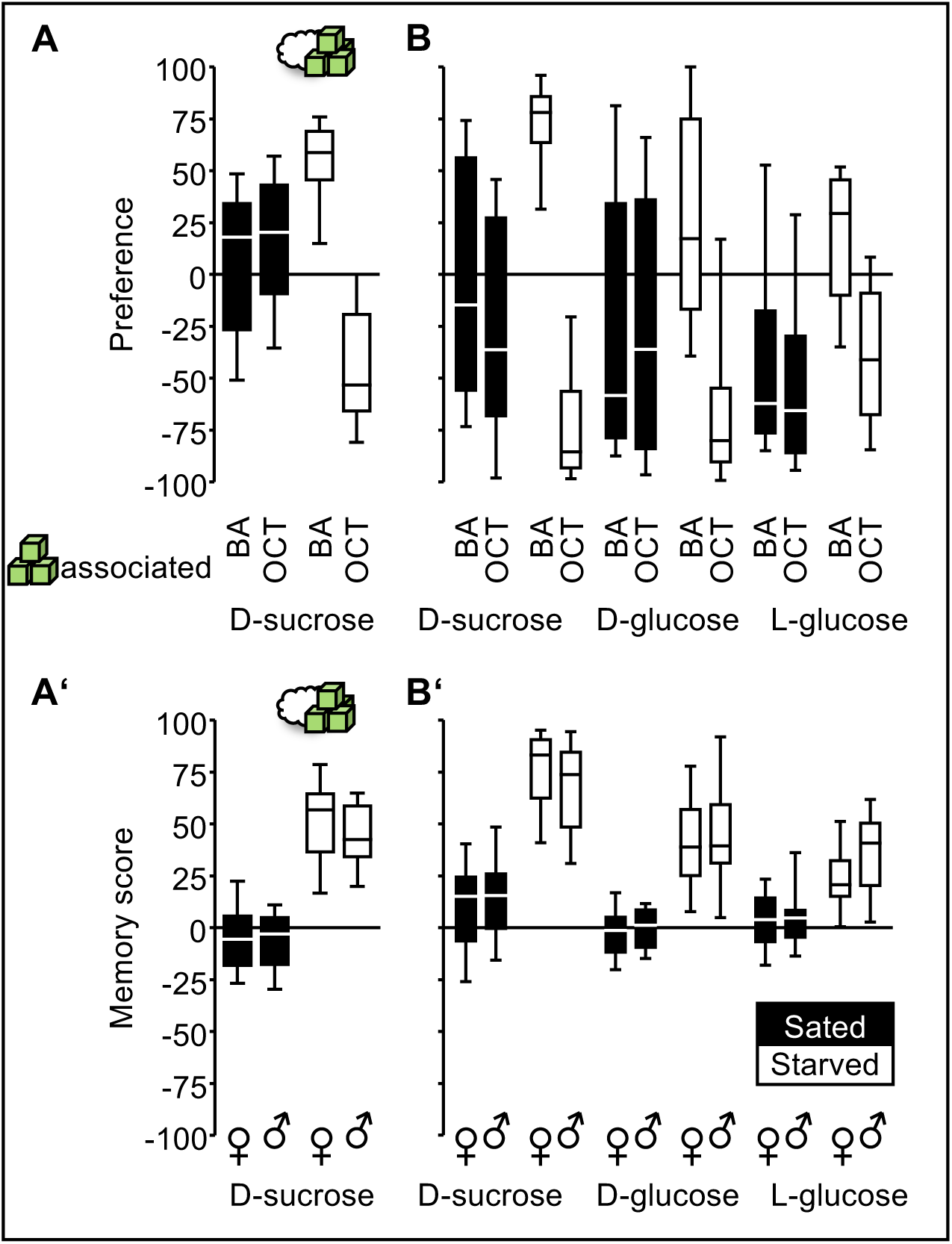
(A) Preference data (see Eqn 1) corresponding to the memory scores presented in Fig. 1B. Filled boxes indicate preference in sated groups, whereas white-filled boxes represent preference in starved groups. The specific odour associated with sugar (BA or OCT) is indicated. **(A’)** Memory scores from Fig. 1B, separated by sex. **(B)** As in (A), for the preference data corresponding to the memory scores presented in Fig. 1C. **(B’)** As in (Á), for the memory scores shown in Fig. 1C. The plotted data are documented in the supplemental data file “Starvation DATA”.

**Fig. S3.**
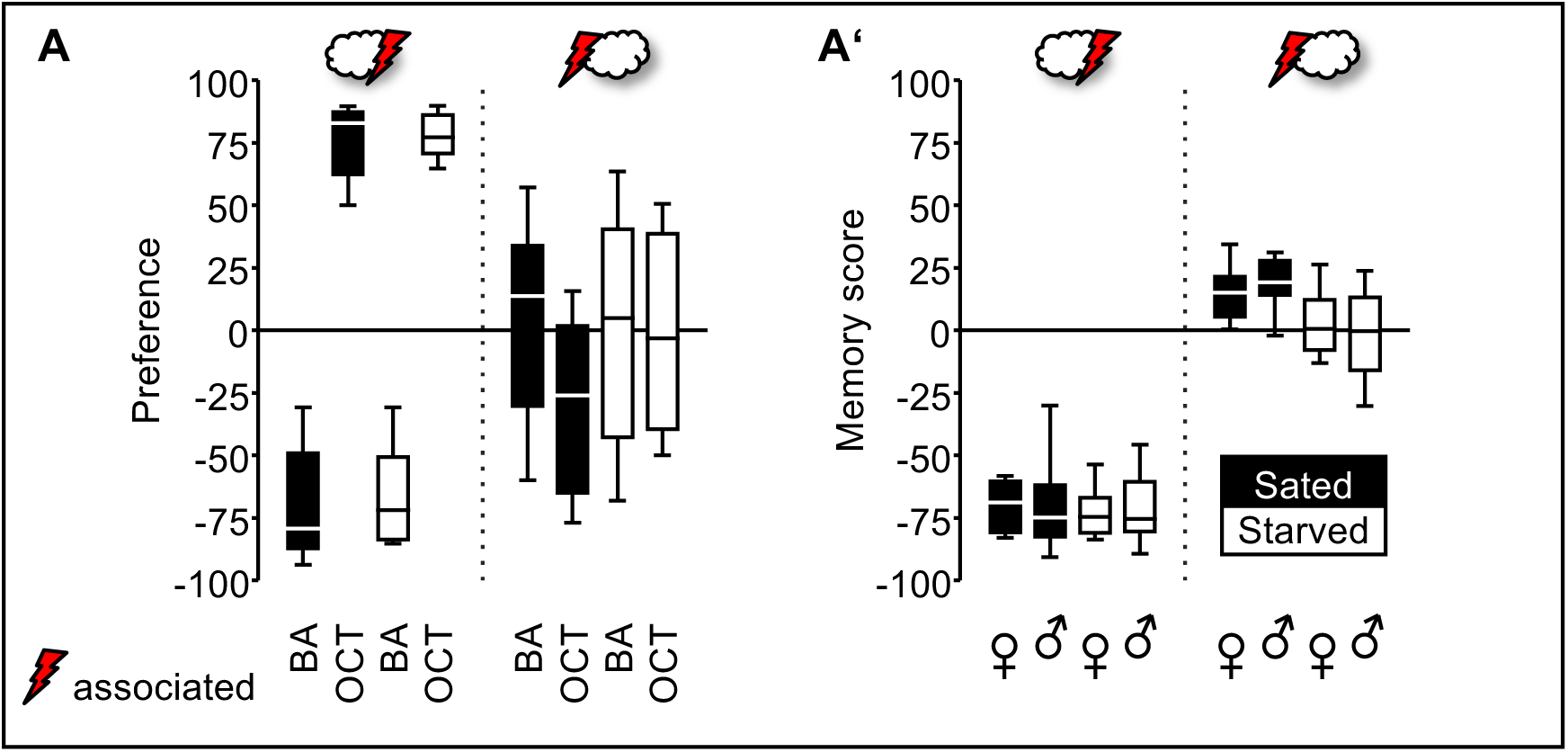
(A) Preference data (see Eqn 1) corresponding to the memory scores presented in Fig. 2B. Filled boxes indicate preference in sated groups, whereas white-filled boxes represent preference in starved groups. The specific odour associated with shock (BA or OCT) is indicated. **(A’)** Memory scores from Fig. 2B, separated by sex. The plotted data are documented in the supplemental data file “Starvation DATA”.

**Fig. S4.**
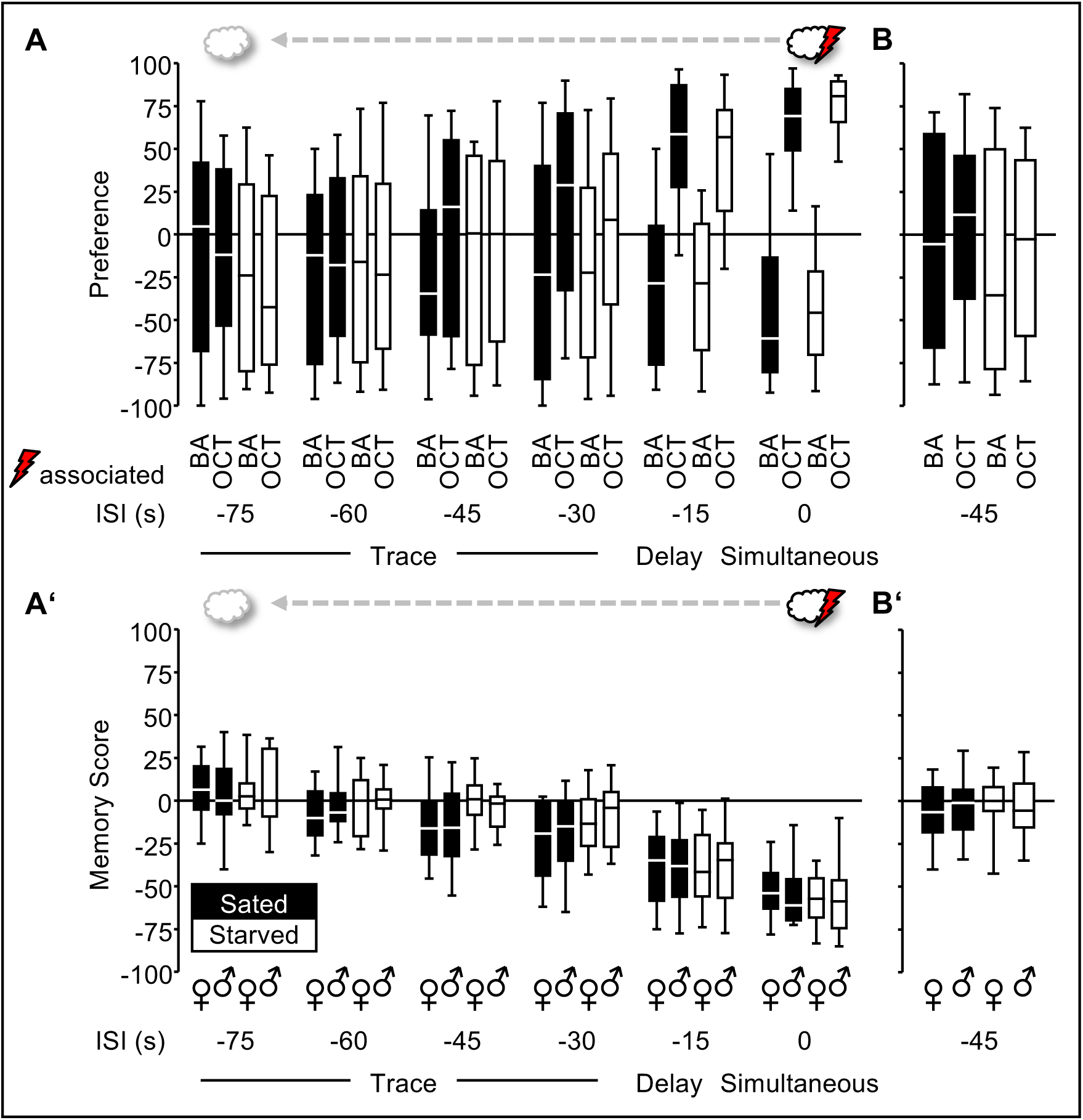
(A) Preference data (see Eqn 1) corresponding to the memory scores presented in Fig. 3B. Filled boxes indicate preference in sated groups, whereas white-filled boxes represent preference in starved groups. The specific odour associated with shock (BA or OCT) is indicated. **(A’)** Memory scores from Fig. 3B, separated by sex. **(B)** As in (A), for the preference data corresponding to the memory scores presented in Fig. 3C. **(B’)** As in (Á), for the memory scores shown in Fig. 3C. The plotted data are documented in the supplemental data file “Starvation DATA”.

**Fig. S5.**
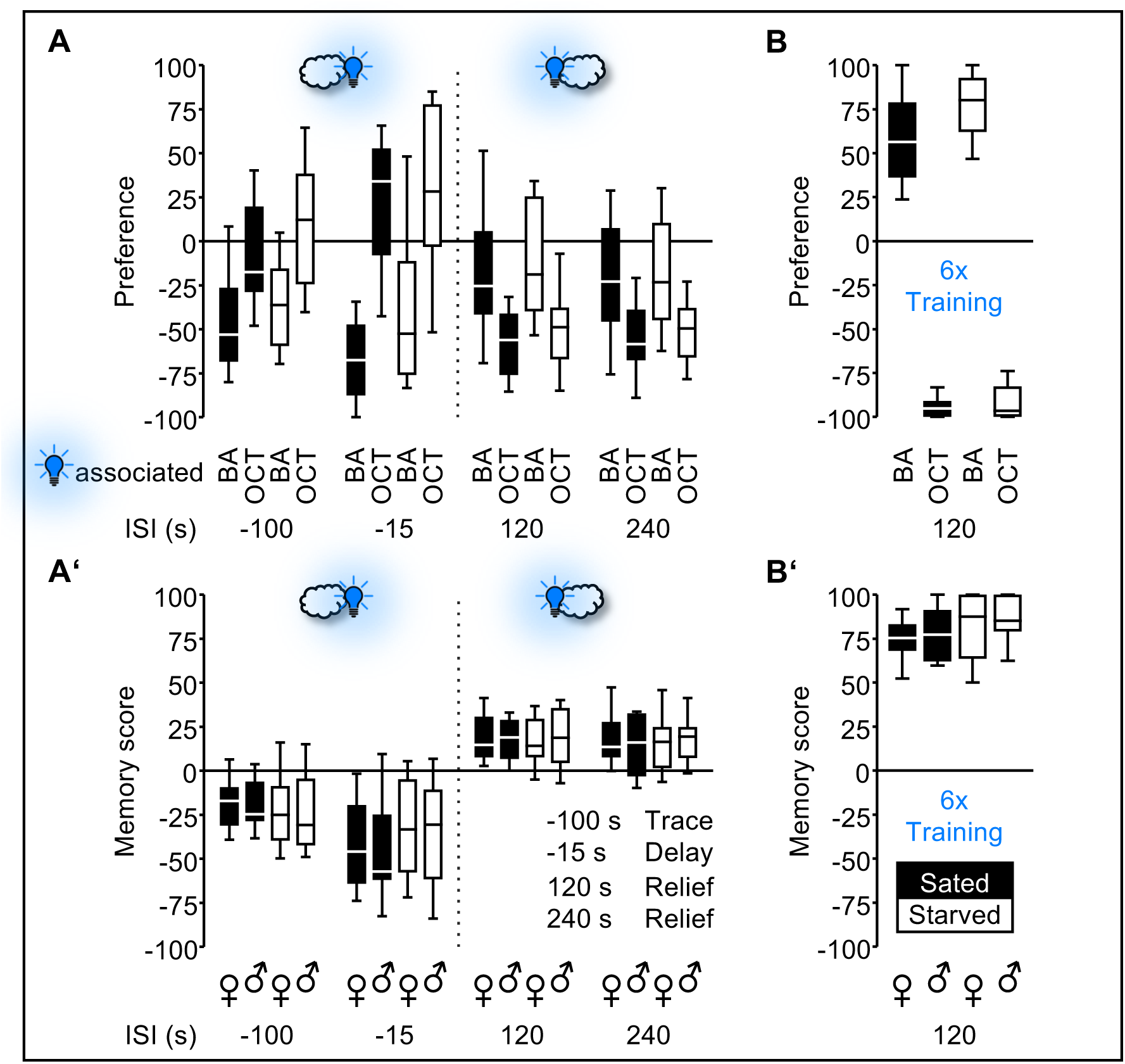
(A) Preference data (see Eqn 1) corresponding to the memory scores presented in Fig. 4C. Filled boxes indicate preference in sated groups, whereas white-filled boxes represent preference in starved groups. The specific odour associated with the optogenetic activation of the PPL1-01 association (BA or OCT) is indicated. **(A’)** Memory scores from Fig. 4C, separated by sex. **(B)** As in (A), for the preference data corresponding to the memory scores presented in Fig. 4D. **(B’)** As in (Á), for the memory scores shown in Fig. 4D. The plotted data are documented in the supplemental data file “Starvation DATA”.

**Fig. S6.**
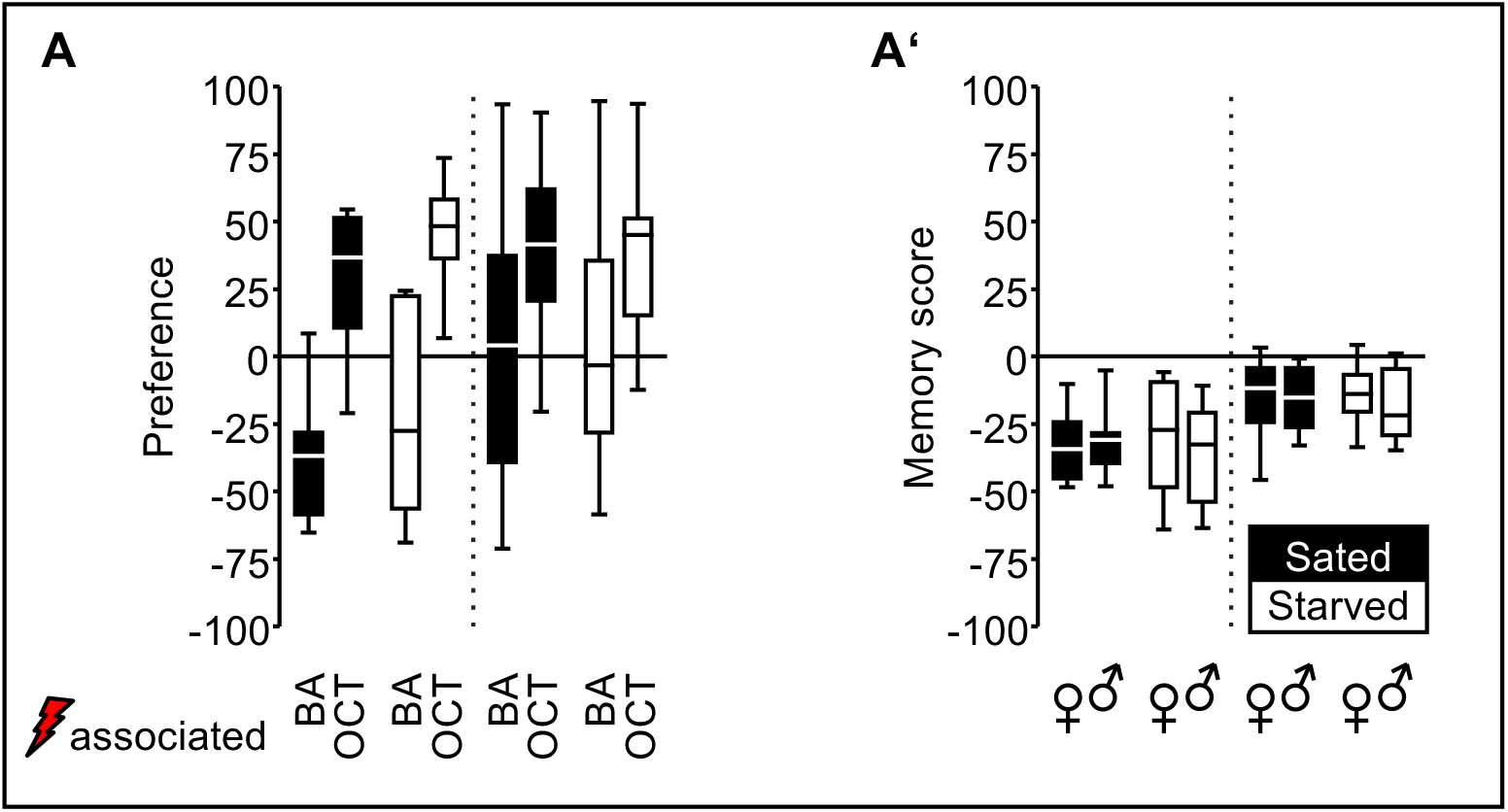
(A) Preference data (see Eqn 1) corresponding to the memory scores presented in Fig. 5B. Filled boxes indicate preference in sated groups, whereas white-filled boxes represent preference in starved groups. The specific odour associated with shock (BA or OCT) is indicated. **(A’)** Memory scores from Fig. 5B, separated by sex. The plotted data are documented in the supplemental data file “Starvation DATA”.

**Fig. S7.**
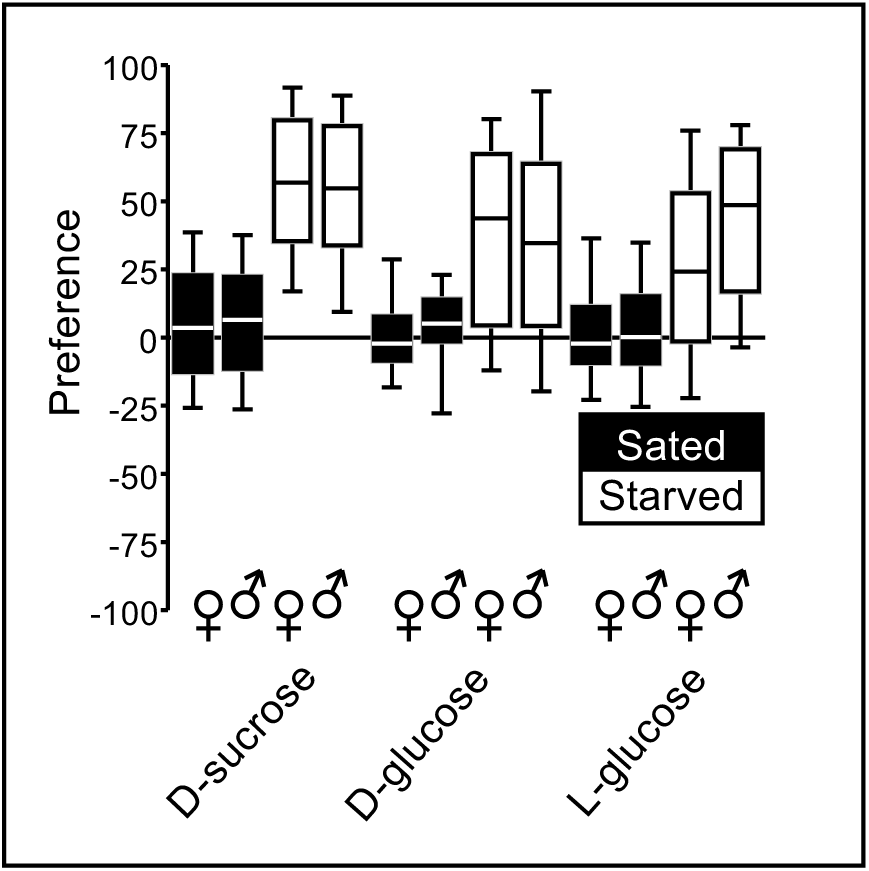
Preference data shown in Fig. 10A, separated by sex. Filled boxes indicate preference in sated groups, whereas white-filled boxes represent preference in starved groups. The plotted data are documented in the supplemental data file “Starvation DATA”.

**Fig. S8.**
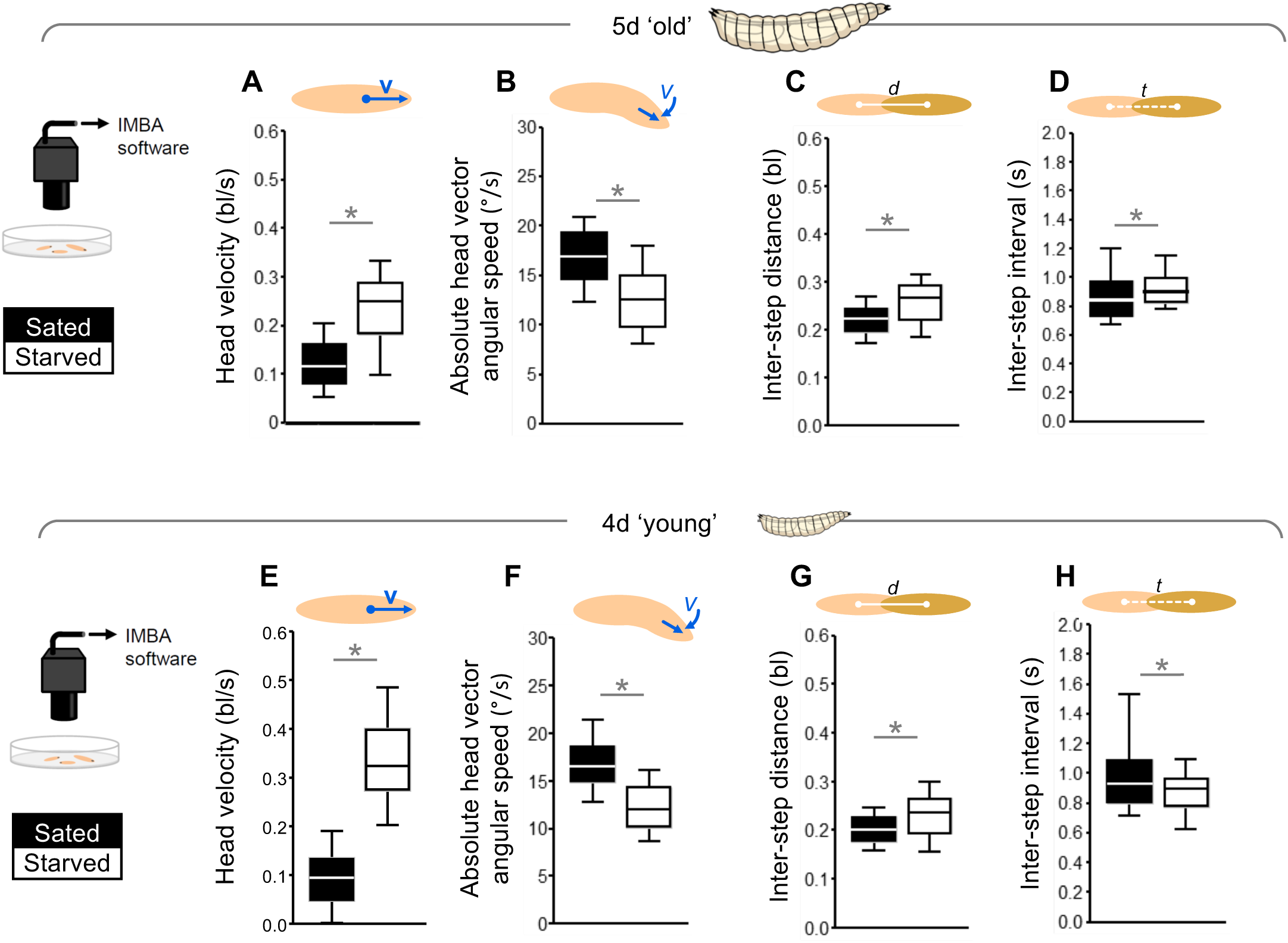
(A-D) Detailed analysis of the data from the experiments shown in Fig. 10F and Fig. 10G for the 5-day ’old’ larvae. Compared to sated larvae, starved animals displayed (A) increased velocity in forward direction (MWU-test, U=2010, p<0.05, N=132, 95), and (B) decreased angular speed (MWU-test, U=2639, p<0.05, N=132, 95) of their heads. Moreover, during peristaltic forward steps, starved animals (C) covered a larger distance (MWU-test, U=3529, p<0.05, N=132, 95), and (D) required slightly more time for each step (MWU-test, U=4942, p<0.05, N=132, 95). **(E-H)** As in (A-D), for the experiments shown in Fig. 10N and Fig. 10O for the 4-day ’young’ larvae. Upon starvation, the larvae displayed the same differences in locomotion as their older counterparts (E: MWU-test, U=544, p<0.05, N=108, 153; F: MWU-test, U=2697, p<0.05, N=108, 153; G: MWU-test, U=5613, p<0.05, N=108, 153) except that in this case the larvae required less time for each step (H: MWU-test, U=6837, p<0.05, N=108, 153). * indicates significance in MWU tests at p<0.05. Statistical results are reported along with the source data in the supplemental data file “Starvation DATA”.

